# Mechanistic Insights on ATP’s role as Hydrotrope

**DOI:** 10.1101/2021.04.28.441722

**Authors:** Susmita Sarkar, Jagannath Mondal

## Abstract

Hydrotropes are small amphiphilic molecules which help in solubilizing hydrophobic entities in aqueous medium. Recent experimental investigation has provided convincing evidences that, adenosine triphosphate (ATP), besides being an *energy currency of cell*, also can act as *hydrotrope* to inhibit the formation of protein condensates. In this work, we have designed computer simulations of prototypical macromolecules in aqueous ATP solution to dissect the molecular mechanism underlying ATP’s newly discovered role as a hydrotrope. The simulation demonstrates that ATP can unfold a single-chain of hydrophobic macromolecule as well as can disrupt the aggregation process of a hydrophobic assembly. Moreover, the introduction of charges in the macromolecule is found to reinforce ATP’s disaggregation effects in a synergistic fashion, a behaviour reminiscent of recent experimental observation of pronounced hydrotropic action of ATP in intrinsically disordered proteins. A molecular analysis indicates that this new-found ability of ATP are ingrained in its propensity of preferential binding to the polymer surface, which gets fortified in presence of charges. The investigation also renders evidence that the key to the ATP’s superior hydrotropic role over chemical hydrotrope (Sodium xylene sulfonate, NaXS) may lie in its inherent self-aggregation propensity. Overall, via employing a bottom-up approach the current investigation provides fresh mechanistic insights into the dual solubilizing and denaturing abilities of ATP.

## Introduction

One of the most important biomolecules of life, Adenosine triphosphate (ATP) serves as the preliminary source of energy in cells to execute important biological processes such as intra-cellular and extra-cellular signalling, neurotransmission, nucleotide and protein synthesis and mainly to drive multiple molecular machines. However, in a recent investigation,Patel and co-workers^1^ experimentally tested the action of ATP on the phase separating proteins such as FUS, TAF15, hnRNPA3, PGL-3 and noticed considerable effect of ATP to solubilize these intrinsically disordered proteins. The work had demonstrated that biological compartmentalisation via liquid-liquid phase separation (of eg. purified fused in sarcoma (FUS)) is dependent on ATP which controls the viscosity^2–4^ of these compartments. As a major implication, the investigation discovered a new role of ATP: apart from being the *energy currency of cell*, it was found to keep the proteins in soluble form and to prevent those from being engaged in any kind of aggregation. In a similar spirit, it was found that high ATP concentration is maintained in metabolically quiescent organ, eye lens, to prevent crowding of *γ*S-crystallin.^5^ Evidences for ATP driven solubilization of aggregates of Xenopus oocyte nucleoli have been reported.^6^ A proteome-wide profiling suggested ATP regulates solubility of a significantly large set of proteins.^7^ These investigations, most of which are recent, have prompted coining a new role of ATP as ‘biological hydrotrope’.^1^ In the present article, we employ a bottom-up approach to probe the ability of ATP to perform as a ‘hydrotrope’ via computer simulation of ATP’s interaction with prototypical hydrophobes and its variants.

‘Hydrotropes’ are the small amphiphilic molecules which help in solubilizing hydrophobic entities in aqueous medium. In 1916, Carl Neuberg^8^ first introduced the term *hydrotropy* for a special class of small organic molecules which can considerably enhance aqueous solubility of very poorly water soluble hydrophobic substances. Likewise, ATP consists of hydrophobic aromatic pyrimidine base ring which is connected via moderately polar five-membered ribose sugar unit to a strongly hydrophilic tetra-negative triphosphate moiety- all these diversified structural units together have built suitable amphiphilic nature in ATP. But how the amphiphilic nature of ATP directly translates to its hydrotropy, which structural part of ATP drives solubilization and most importantly the underlying molecular principle are currently elusive. The intricate mechanism involving ATP’s hydrotropy and specifically the combination of the precise interactions of ATP with the macromolecules it is attempting to solubilize, is not completely explored. Since ATP, in its newly ordained role as ‘hydrotrope’, can potentially play promising role in several aspects of biology there have been multiple attempts to evaluate the concerned mechanism. He et al. suggested that ATP antagonises the crowding by resisting the overlap of the hydration shells around different solute molecules but without any direct interaction with the solubilizate. ^5^ This investigation implied that the disaggregating ability of ATP is not resulted from the clustering of ATP molecules around a hydrophobic patch. On the other hand, Patel et al.^1^ proposed that the hydrotropic action of ATP proceeds through formation of dynamic clusters over a solute particle. These contrasting hypotheses on ATP’s hydrotropy preclude positing an unanimous mechanism at play here. Again while Kang et al. ^9^ emphasizes only pi-cation and electrostatic interactions as the driving forces behind ATP’s hydrotropic role, the proposal by Pal et al. ^10^ came up with relative contributions from van der waals and electrostatic interactions, H-bonding, *π*−*π* and NH-*π* interactions guiding ATP’s hydrotropy in a combinatorial manner. Therefore a clear picture is yet to be emerged.

In fact the generic interpretation of hydrotropy lacks clarity. In 1946, McKee interpreted hydrotropy as a salting-in process. ^11^ However, according to Winsor et al, hydrotrope acts as a common solvent for the hydrophilic and lipophilic compounds, ^12^ suggesting hydrotropy as a process similar to co-solvency. In 1966, Ueda pointed out the formation of hydrotropic complex with solubilizate and thus the decreased activity coefficient in water, are responsible for enhanced solubility by hydrotrope. ^13^ On the other hand, Rath was concerned about the hydrotropic structural features and its stacking around the solute as the effective reason towards hydrotropy^14^. But according to Balasubramanian et al. ^15^ the cooperativity adopted by hydrotrope molecules is low. Thus a precise understanding of the hydrotropic mechanism has been elusive over the years. ATP’s recent emergence as a biologically occurring hydrotrope has added to the existing puzzle.

In the current investigation, we aim to identify the physical principles innate to the hydrotropic mechanism of ATP via computer simulation. Here we employ Molecular Dynamics (MD) simulation for exploring the role of ATP as intracellular solubilizer and as well as natural denaturant. Instead of grappling with the complexity associated with proteins or biopolymers, we computationally design a prototypical scenario of a bead-in-a-spring macromolecule solvated in atomically modelled aqueous solution of ATP via classical force fields. The simple and bottom-up nature of the model of macromolecules allow us the flexibility needed for tuning the interactions and introducing charges on them. We investigate the action of ATP on 1) conformational landscape of single macromolecule and 2) the aggregation propensity of the multiple copies of same chain. Our investigation uncovers the denaturing and solubilizing ability of ATP towards hydrophobic macromolecules, a signature of hydrotrope. We also determine how the presence of charge and different charge distributions would influence ATP’s role as hydrotrope. The investigations combine Wyman-Tanford preferential interaction theory^16,17^ and free energy calculation (umbrella sampling method) ^18^ to elucidate the underlying mechanism of ATP’s hydrotropic action. Finally, we demonstrate ATP’s superior efficacy over chemical hydrotrope Sodium xylene sulfonate (NaXS) and hyopthesize the key lies in ATP’s inherent ability to self-assemble.

## Materials and methods

### Simulation model

Single and multiple copies of a prototypical 32-bead hydrophobic polymer chain, solvated in aqueous ATP solution, form the basis of the current investigation. This polymer has been extensively studied by Mondal et al. in past and the details of interactions associated with the polymer chain can be found elsewhere. ^19^ All weak dispersion nonbonding interactions have been considered as Lennard Jones (LJ) interactions (*σ* = 0.4 nm and *ϵ_p_* = 1.0 kJ/mol). Additionally, to investigate the influence of coulomb charge, in separate investigations, we have selectively assigned charges on certain polymer beads, resulting in design of four additional polymers: ‘purely negative polymer’ (4 individual negative charges are placed periodically), ‘purely positive charged polymer’ (4 individual positively charges are placed periodically), ‘alternative charge-neutral polymer’ (4 positive and 4 negative charges are placed alternatively throughout the chain) and ‘block charge-neutral polymer’ (like charges are placed in one block and the opposite like charges are placed in other block of the polymer). Figure 1 depicts the topology of respective polymer chains decorated with various charge-patterns on selected beads. In all of the cases, the partial charge of the individual charged beads are maintained at |*q*| = 1e. The charges on the polymer are modelled by Coulomb interaction. We refer to figure 1 for chemical structures of cosolutes (ATP or NaXS) and solvent (water) employed in the work. Water molecules are modelled using charmm-TIP3P interaction potential,^20^ while CHARMM36 force field^21^ is applied for ATP.

**Figure 1:**
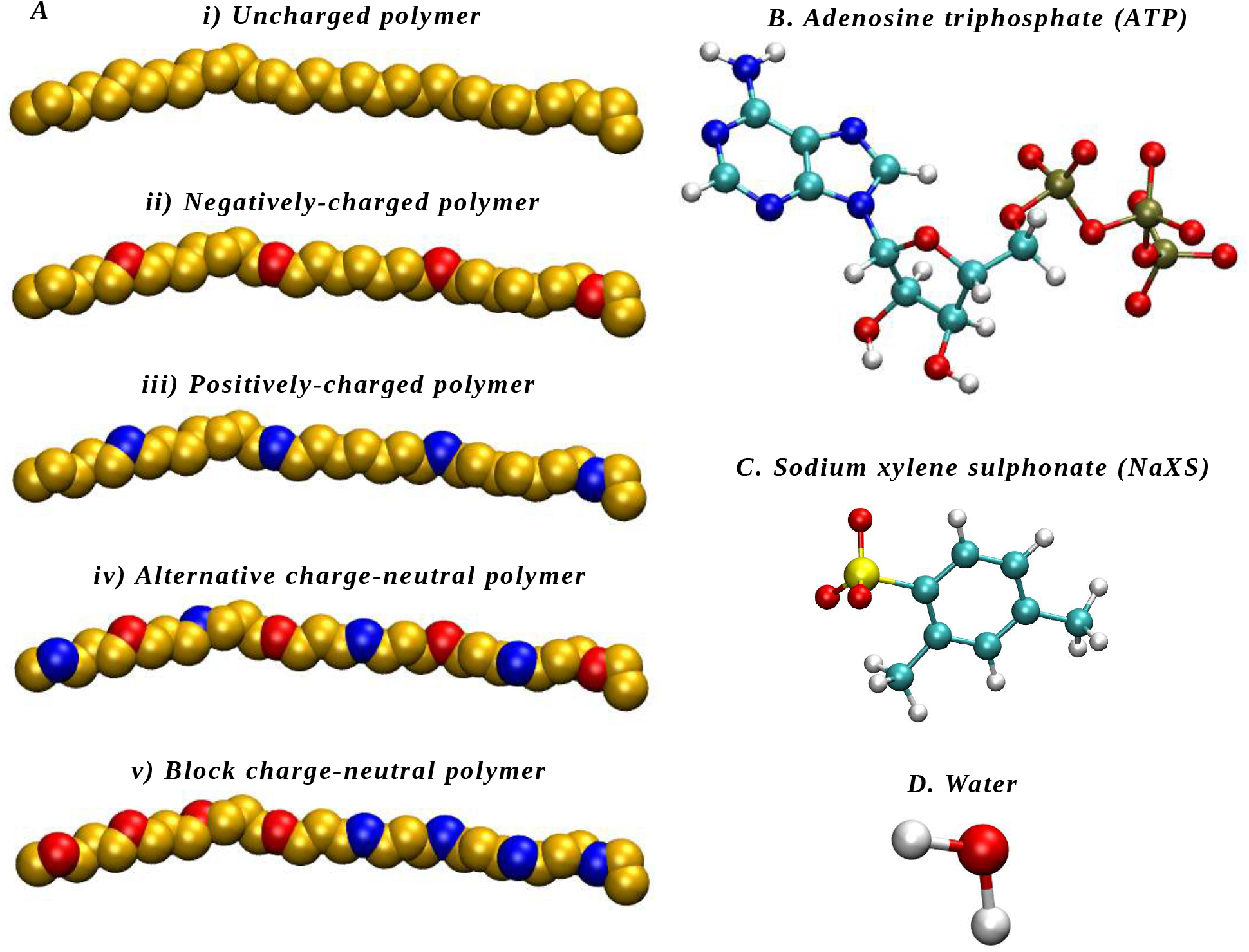
A. Representative extended configurations of polymer with a different number and distribution of charged beads. Red, blue and golden colors represent positive, negative and no charge respectively.(i)uncharged,(ii)negatively charged,(iii)positively charged,(iv)alternative charge-neutral and (v) block charge-neutral polymer. B. A representative ATP molecule C. A representative NaXS molecule D. A representative water molecule

We have also compared the efficiency of ‘biological hydrotrope’ ATP against a ‘chemical hydrotrope’ NaXS (previously investigated in experiments by Patel et al). The CHARMM generic force field (CGENFF)^22^ parameter of NaXS was re-optimised against ab-initio charge and dihedral scans via ‘GAAMP’ protocol.^23^ The charge of ATP is neutralized by Mg^2+^ ion (parameterised using Charmm36 force field) and for neutralization of the negative charge of the negatively charged polymer, Mg^2+^ ion is used whereas in case of positively charged polymer Cl^−^ ion is inserted for neutralization and in case of NaXS, system is neutralized by Na^+^ ion.

#### Simulation Method

We have performed classical molecular dynamics (MD) simulations to investigate a) conformational landscape of a single copy and b) aggregation propensity of multiple copies of the 32-bead polymer chain and its variants, solvated in variously concentrated aqueous solution of ATP. Tables T-1 and T-2 in Figure 2 provides the details of two main types of computer simulations employed in the current work:

**Figure 2:**
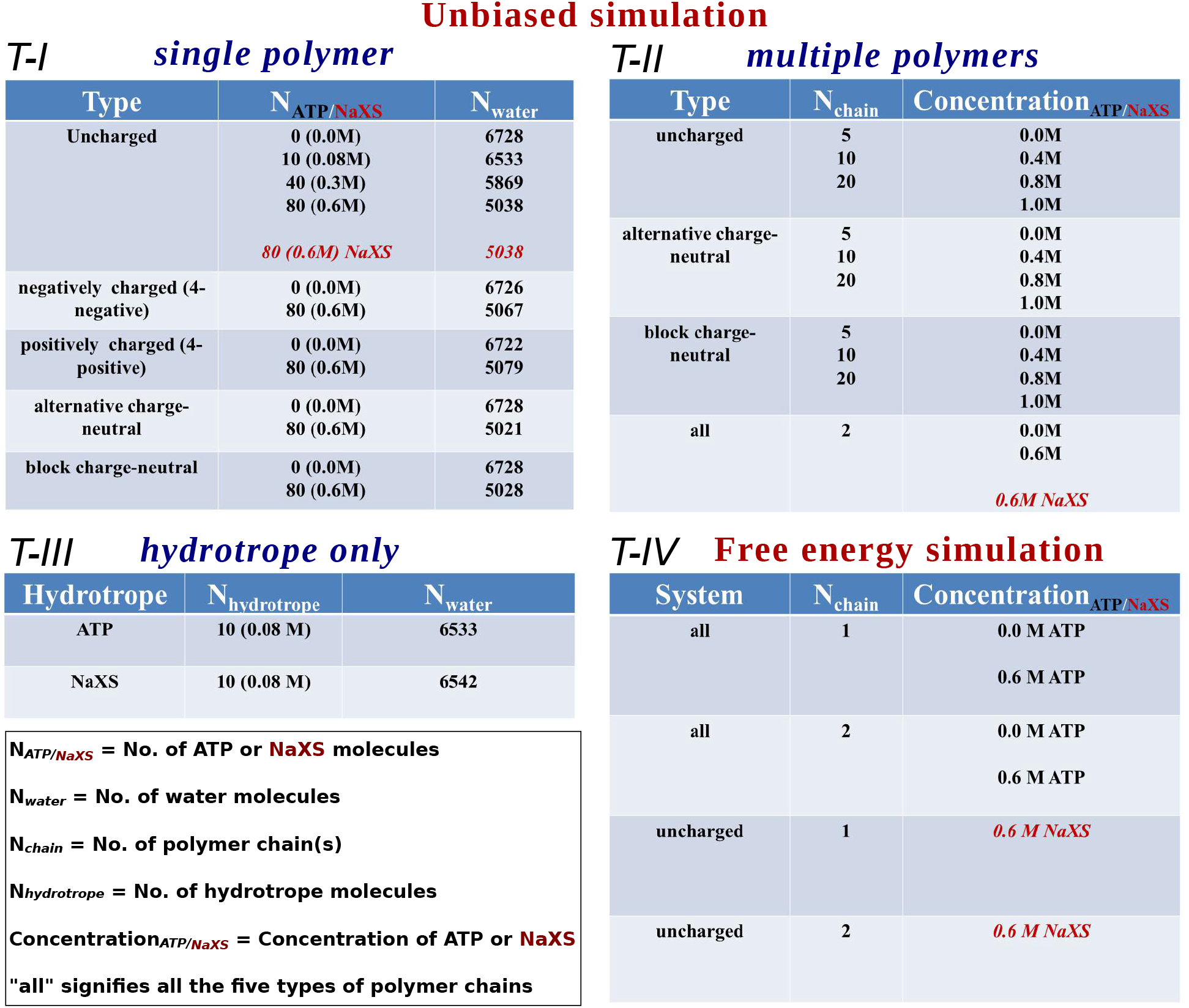
Table I-III represents the details of unbiased simulation with single polymers, multiple polymers and only hydrotrope molecules respectively. Table IV is for the presentation of the simulation details of the entire free energy simulation with single and double polymer chains in presence of ATP and NaXS as hydrotrope

As enumerated in Table T-1, we have performed equilibrium MD simulations with single copy of the uncharged hydrophobic polymer in a wide set of aqueous ATP solutions to investigate the effect of ATP on conformational landscape. The simulated media consisted of neat water (i.e. 0 M), 0.08 M, 0.3 M and 0.6 M aqueous ATP solution (see T-1 in figure 2 for system details). To investigate the effect of charge in polymer on the ATP-induced conformational landscape, each of the charged polymer listed in figure 1 was simulated in 0.6 M ATP solution (and compared with that in neat water). Each MD simulation trajectory was 100 ns long in which the polymer exhaustively sampled multiple key conformations. Each of the simulations were replicated five times for assessing the statistical robustness. All the simulations have been performed within a cubic box of dimension 5.86 nm in each direction.

For investigating the solubilizing ability of ATP as a hydrotrope, we simulated the aggregation propensity of multiple polymer chains in a wide range of concentration of ATP. Table T-II in figure 2 provides the number of polymer copies simulated for investigation of self-aggregation propensity in various concentrations of the ATP solution. Specifically, we have simulated individual cases of multiple polymeric systems consisting of 5, 10 and 20 uncharged polymeric chains in 0 M, 0.4 M, 0.8 M and 1 M of aqueous ATP solutions. The same is repeated for exploring aggregation propensity in ‘block charge-neutral’ and ‘alternating charge-neutral’ polymer systems (see figure 1 for the topology). All systems are simulated for 500 ns and to avoid biases, each of the uncharged multiple polymeric simulations were replicated seven times. For each of the simulation the cubic box size is kept constant at 5.86 ×5.86 ×5.86nm^3^. The aggregation propensity is characterised via visual inspection as well as cluster size distributions. The cluster was ascertained if the distance between the centre of mass of a pair of polymer chains is within 0.5 nm.

Each of the systems consisting of single polymer chain in aqueous ATP solution is first subjected to energy minimisation via steepest-descent algorithm followed by final 100 ns MD simulation in NPT ensemble at an average temperature of 303 K- maintained by velocity rescale thermostat (time constant *τ_T_* = 1.0ps) and at equilibrium pressure of 1bar- managed with Parrinello-Rahman barostat^24^ (time constant *τ_P_* = 2.0ps). Each simulation is performed in cubic box of length 5.86nm. For the simulation with multiple polymers, after energy minimisation, 500 ps NVT equilibration is carried out followed by 500 ns final production run in NPT ensemble keeping temperature and pressure at 303K and 1bar maintained by the above mentioned temperature and pressure coupling schemes with the similar time constants as before. Perioidic boundary condition is implemented in all three dimensions. A leap-frog integrator with a time step of 2 femtosecond was employed for time propagation. A cut-off of 1.2 nm has been chosen in case of van der Waals and Coulomb interaction. Particle-Mesh Ewald (PME)^25^ method with a grid spacing of 0.12 nm has been employed for treatment of long-range electrostatic interactions. To constrain the bonds associated with hydrogen and for bonds as well as angle of water molecules, LINCS^26^ and SETTLE^27^ algorithms respectively are employed in our simulation. For Energy and Pressure, long range dispersion corrections has been used. 2019.3, 2018.6 and 5.0.6 version of GROMACS software ^28,29^ are employed to execute all the molecular dynamics (MD) simulations.

Apart from the investigation via equilibrium MD simulation, we have also characterised a free energetic perspective of the ATP-induced modulation of conformational landscape and aggregation tendency of the macromolecule(s). Table T-IV of figure 2 provides the system details employed for umbrella sampling simulations. Towards this end, we have employed umbrella sampling simulations along radius of gyration (Rg) of the single chain polymer (for all topologies as enumerated topology in figure 1) in neat water and that in 0.6 M aqueous ATP concentration, using the same protocol as in the previous work of Mondal et al. ^19^ We found harmonic restraints of force constants ranging between 3500-5000 kJ/mol/nm^2^ at windows corresponding to Rg ranging between 0.4 nm to 1.2 nm at a regular interval of 0.05 nm were optimum in exploring the desired windows and allowing sufficient overlaps between the adjacent windows. Each of the windows for umbrella sampling simulation are sampled in the NPT ensemble for 20 ns. We have implemented PLUMED extension^30,31^ of gromacs for free energy simulation. The box size is same in case of each of umbrella sampling simulation as reported for unbiased simulation.

Like-wise, to quantify the ATP-induced aggregation process, the equilibrium MD simulation was supplemented with the free energy calculation of aggregation of a pair of the polymer copies in neat water and in 0.6 M aqueous ATP concentration. In this case, the distance between centre of mass of the polymers (*d*) was employed as a collective variable for umbrella sampling, with the value of *d* varying between 0.1 nm to 1.5 nm at a spacing of 0.1 nm. We found that the employment of a harmonic restraints force constants ranging between 3500-5000 kJ/mol/nm^2^ over the windows or sometimes application of only one of those to all the windows corresponding to a particular biased simulation was optimum for maintaining sufficient overlap among adjacent windows. In this case also the simulation box size is kept as it has been used earlier.

The weighted histogram analysis method (WHAM)^32^ has been used to calculate the unbiased histograms and the corresponding free energies by combining the data of each independent trajectories for each window. Each of these free energies corresponding to both the individual collective variables (Rg for single polymer and d for a pair-of-polymer system) has been statistically averaged over 2 independent trajectories and the standard deviations have been represented as error bars.

The differences in free energy (Δ*G^CE^* = *G^E^* − *G^C^* and Δ*G^AS^* = *G^S^* − *G^A^*) between the extended (E) and collapsed (C) conformations of the single polymer and the separated (S) and assembled (A) state of the two polymer(s) has been compared across different solutions (0M and 0.6M) without and with ATP for each of the polymer variants described in figure 1. We have specifically calculated the change of Δ*G^CE^* and Δ*G^AS^* (hereby defined as ΔΔG) in the aqueous ATP solution relative to that in neat water with the aim of a systematic comparison across all different polymer chains and to discard the intrinsic effect of the nature of polymer from that of neat water.

ΔΔG is expressed as:

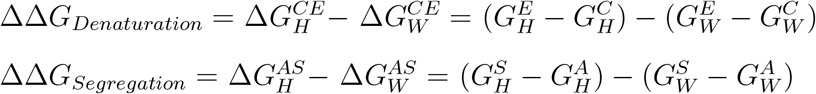

where H refers the binary mixture containing hydrotrope-ATP and solvent-water and W stands for neat water situation.

We have performed similar umbrella sampling simulation with the single uncharged polymer for NaXS (0.6 M) in order to compare the efficiency of ATP relative to NaXS in denaturing a polymer chain. For a comparative study of the ability of hydrotrope driven segregation, we have designed the similar set of umbrella sampling simulation containing double uncharged polymer chains in aqueous solution of NaXS (0.6 M). We have done equivalent equilibrium simulation (following similar protocol as before) with only single and a pair of uncharged polymer(s) (as control) in 0.6M NaXS solution for 100 ns with an aim to produce representative conformations for umbrella sampling simulation.

In addition to check the ability of self aggregating tendency among hydrotropes we have simulated 10 ATP and NaXS molecules separately solvated with water (0.08 M concentration), in NPT ensemble for 100 ns. For all the simulations cubic box with a dimension of 5.86 nm in each direction was considered.

## Results and Discussion

### ATP promotes unfolding of single-chain hydrophobic macromolecule

We first investigate the effect of ATP concentration on the conformation of a hydrophobic polymer chain. Towards this end, we use radius of gyration of the polymer (*R_g_*) as a collective variable of polymer size. Figure 3 A shows the representative time-profiles of *R_g_* of the polymer in multiple ATP solutions. Starting with an extended conformation (*R_g_*=1.2 nm), the polymer folds fairly quickly to a collapsed conformation (*R_g_*=0.45 nm) in neat water. This reproduces past observations by Mondal et al^19^ for the same polymer. Interestingly the current simulation also reveals that gradual incorporation of ATP in aqueous media arrests the polymer collapse to a relatively higher *R_g_* value than that sampled in neat water. As evident from figure 3, in 0.08 M and 0.3 M aqueous ATP solution, the hydrophobic polymer attains a conformation corresponding to average *R_g_*=0.6 nm, which is representative of a hair-pin like structure. On the other hand, at a high concentration of 0.6 M ATP, the polymer remains extended at a conformation corresponding to *R_g_*=1.0 nm, suggestive of ATP’s role in preventing the folding/collapse behaviour of a macromolecule. The overall observation is statistically reproducible across multiple independently performed unbiased MD trajectories. (See Figure S1).

**Figure 3:**
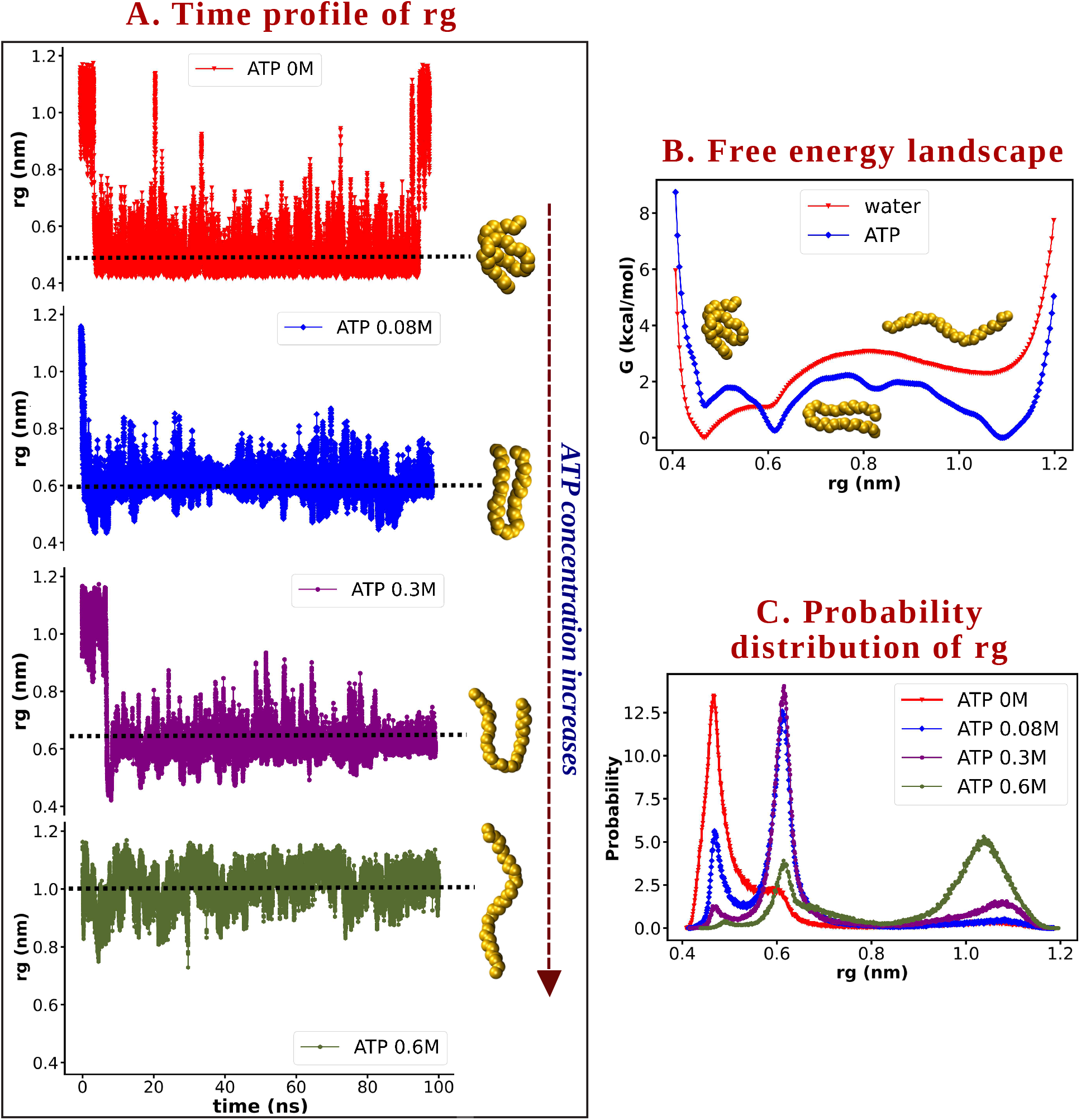
Single chain hydrophobic polymer: A. Plot of time profile of *R_g_* of uncharged hydrophobic polymer (see figure 1A i) in 0.0M ATP (red), 0.08M ATP (blue), 0.3M ATP (violet) and 0.6M (olive green). B. Free energy landscape of single polymer chain in neat water (red) and in 0.6M aqueous solution of ATP (blue). Representative folded, semi-folded and un-folded configurations of the model 32-bead polymer are shown in the corresponding rg value represented in the figure. C. Probability distribution of the Rg of the uncharged single polymer chain in multiple different concentrated solution of ATP (0M, 0.08M, 0.3M and 0.6M) in water.

For a more quantitative characterisation of ATP’s role in modulating the conformational landscape of the macromolecule, we compute the free energy profile of the polymer as a function of *R_g_* via umbrella sampling (see Method). Figure 3B compares the free energy profile of polymer in 0.6 M aqueous ATP solution with that in water. The free energy landscapes both in neat water and aqueous ATP solution indicate three basins corresponding to collapsed (C) (*R_g_*=0.45 nm), hairpin (*R_g_*=0.6 nm) and extended (*R_g_*=1.0 nm) (E) conformation. However, as depicted by the free energy profile in figure 3B, polymer’s extended conformation is found to be free energetically stabilised, while collapsed conformation is destabilised in aqueous ATP solution. The systematic trend of unfolding ability by ATP is evident in the increased probability of occurrences of conformation (figure 3C) corresponding to higher *R_g_* of the polymer. Overall, the simulation results in figure 3 indicate that ATP is discouraging the polymer to be in folded conformation and thereby favouring denaturation of the polymer chain.

### ATP solubilizes hydrophobic aggregates

The previous discussion provided clear evidence for ATP’s role in preventing hydrophobic polymer collapse at a single chain level and promoting extended conformation as a denaturant. The observation prompted us to explore how ATP would influence the aggregation of similar hydrophobic polymers. This is an important question which can directly assess ATP’s ability to sollubilize an aggregate, the hall-mark action of a typical ‘hydrotrope’. Towards this end, we individually simulate the aggregation process of (see Materials and methods) 5, 10 and 20 randomly dispersed hydrophobic polymers in four different aqueous ATP solutions, i.e. 0M, 0.4M, 0.8M and 1M. Figure 4 illustrates the overall results of our investigation of aggregation process in diverse ATP concentration. Figure 4A depicts that, in our MD simulation, which were initiated with 10 randomly dispersed chains, the polymers coalesce into a single large aggregate in neat water. On the other hand, in aqueous ATP solution the representative snapshot indicates presence of multiple aggregates of relatively smaller size than a single aggregate, indicating that ATP solubilizes the single aggregate of hydrophobic polymers into multiple entity. The radial distribution function (RDF) among the polymers (figure S2) shows that with increasing concentration of ATP in the system, the probability of the polymers to be mutually closer decreases. A comparison of polymer’s cluster-size distribution in a representative (0.8 M) ATP solution with that in neat water confirms ATP’s sollubilizing ability of hydrophobic aggregate: in both systems containing 10 and 20 copies of polymer chain, introduction of ATP results in multiple smaller aggregates than a single large aggregate (Figure 4B).

**Figure 4:**
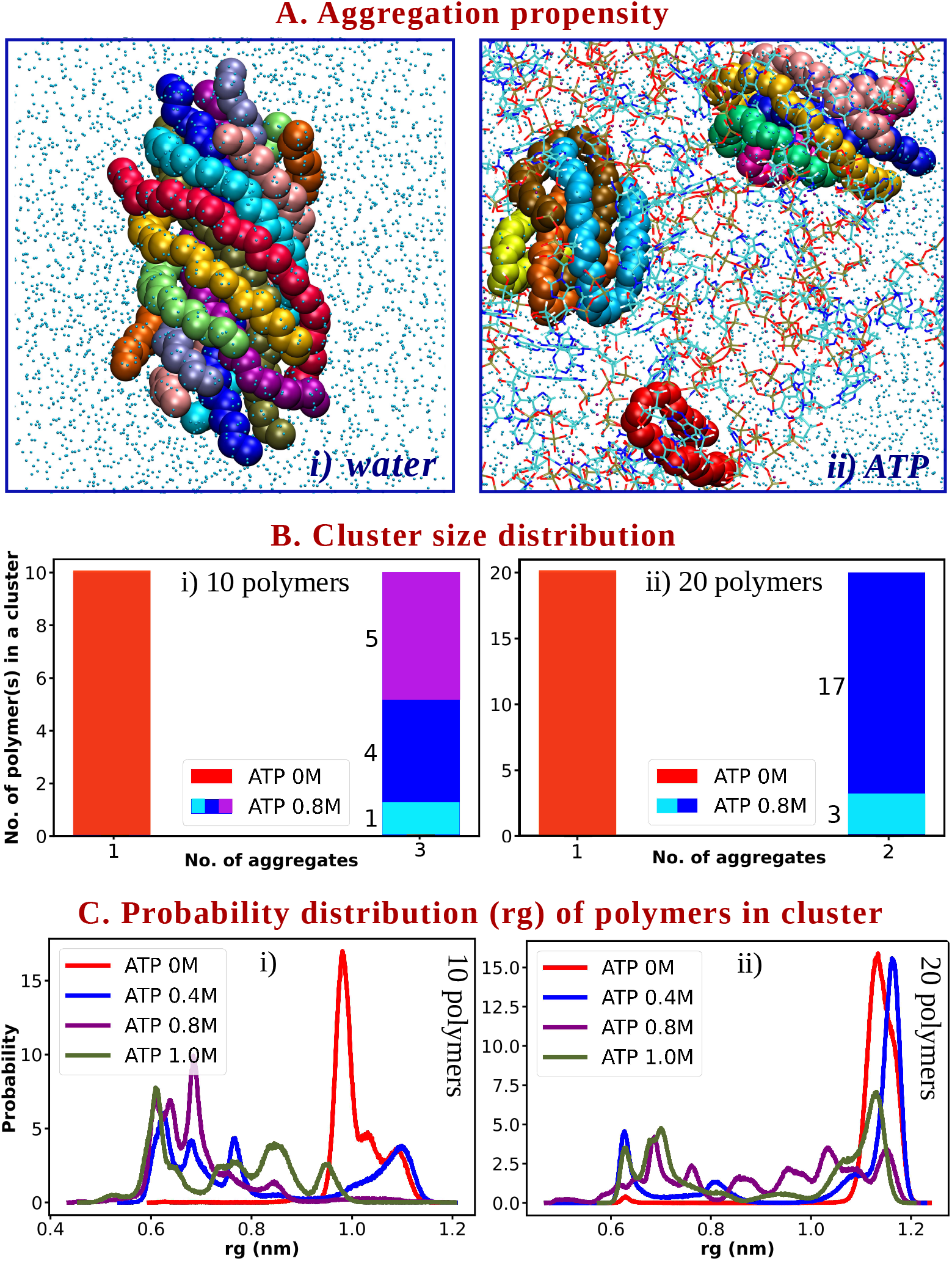
Aggregation of hydrophobic polymer chains: A. Representative snapshots showing aggregation propensity of 10 copies of hydrophobic polymer chains in i) neat water and in ii) 0.8M aqueous ATP solution. In neat water all the 10 polymer chains have formed one single aggregate where-as in 0.8M ATP solution 3 different smaller aggregates exist consisting 1, 4 and 5 polymers B. The bar plots are showing the cluster-size distribution of i) 10 and ii) 20 polymeric systems at 0.8M ATP solution with respect to neat water. In absence of ATP both the polymeric systems (with 10 and 20 polymer chains) forms one single aggregate (red) consisting all the polymers present in the system (10 and 20 respectively). But in 0.8 ATP solution for 10 polymeric system 3 individual aggregates are formed consisting of 1 (light blue), 4 (blue) and 5 (violet) chains. In case 20 polymer chains in 0.8M ATP medium, 2 aggregates with 3 (light blue) and 17 (dark blue) polymer chains are observed. C. Probability distribution of *R_g_* of individual polymer chains involved in aggregate for i) 10 and ii) 20 polymer systems respectively in all the four concentrations of ATP 0.0M (red), 0.4M (blue), 0.8M (viiolet) and 1M (olive green).

Interestingly, upon visual inspection, the individual polymers in an aggregate (see figure 4A), are all found to be in extended conformation in non-ATP medium. This might apparently be in discord with their preferred single-chain conformation in neat water (as was presented in figure 3). This is further characterised by predominantly single peak in probability distribution at around *R_g_*=1.0 nm in neat water (see figure 4C, red curve). This observation can be rationalised by the fact that multiple chains, when present in an ensemble in neat water, tend to maximise the surface area of hydrophobic interaction via remaining extended. The possibility of favorable interaction among the extended uncharged polymers via hydrophobic interaction, can potentially counterbalance the penalty due to exposure of the unfolded surfaces in pure water. Thus these chains can easily be involved in the aggregation and will eventually loose the solubility in water. On the other hand, in an aqueous ATP media, the large aggregate can not be formed as before. ATP molecules tend to tune the radius of gyration (figure 4C) of the polymers so as to decrease the extent of hydrophobic interaction among them such that the formation of large aggregate can be averted and polymers can only be involved in smaller aggregates (sometimes with the shape of ‘james clip’ consisting of hairpin polymers). Taken together, the hydrophobic interaction among the uncharged polymers get disrupted in presence of ATP, which would not allow all the polymers to be engaged in a single aggregate and thereby ATP could favor disaggregation of hydrophobic entities.

In summary, from the study of the single and multiple uncharged hydrophobic polymeric systems, it is evident that there are two prominent roles of ATP-1) denaturation of the single hydrophobic chain and 2) segregation of hydrophobic entities, similar to a hydrotrope.

### Presence of charge in macromolecule reinforces ATP’s unfolding Ability

The discussion till now has confirmed that the presence of ATP in aqueous solution would assist in unfolding of single hydrophobic chain and dissolution of hydrophobic aggregate.

This triggers an immediate question: how does the induction of ATP’s hydrotropy depend on the chemical nature of the macromolecule of our interest? To answer this question we design a series of polymers with diverse charge pattern and first investigate ATP’s ability in modulating the conformational landscape of single copy of these polymer chains. Figure 1A depicts the snapshot of the polymer chains employed in this work: These are coined as ‘uncharged’, ‘negatively charged’, ‘positively charged’, ‘alternative charge-neutral’ and ‘block charge-neutral’ polymer in the current work. Figure 5B shows free energy profiles of the ‘block charge-neutral’ polymer in neat water and in 0.6 M aqueous ATP solution, computed using umbrella sampling approach. The free energy difference indicates that, relative to neat water, ATP solution stabilises the extended conformation of this polymer and destabilises the collapsed conformation. Figure S3 compares the free energy landscapes of all five different polymer chains in presence of ATP with that in absence of ATP. We find that, the qualitative trend that ATP stabilises the extended conformation relative to the collapsed conformation remains robust for all types of macromolecules employed here. However, as quantified by the ΔΔ*G* values in figure 5A, extent of stabilisation depends on the presence, type and decoration of charge. A negative value of ΔΔ*G* in all cases unambiguously confirms that irrespective of the nature of the polymeric chain the process of denaturation by ATP is favourable for all the cases. Nonetheless the comparison of ΔΔ*G* also indicates that ATP’s effect becomes more pronounced with the presence of charge in the polymer. It is observed that ΔΔ*G* value of unfolding gradually increases from uncharged to charge-neutral polymer through the net charged polymer. Thus it can be concluded that electrostatic charge reinforces ATP’s ability to denature polymer chain. Moreover, relatively more negative value of ΔΔ*G* for charge-neutral system compared to the negatively or positively charged polymers suggests that the presence of both the charges (positive and negative) along with uncharged beads amplifies ATP’s effect to a greater extent than the presence of either negative or positive charge with uncharged beads in the system.

**Figure 5:**
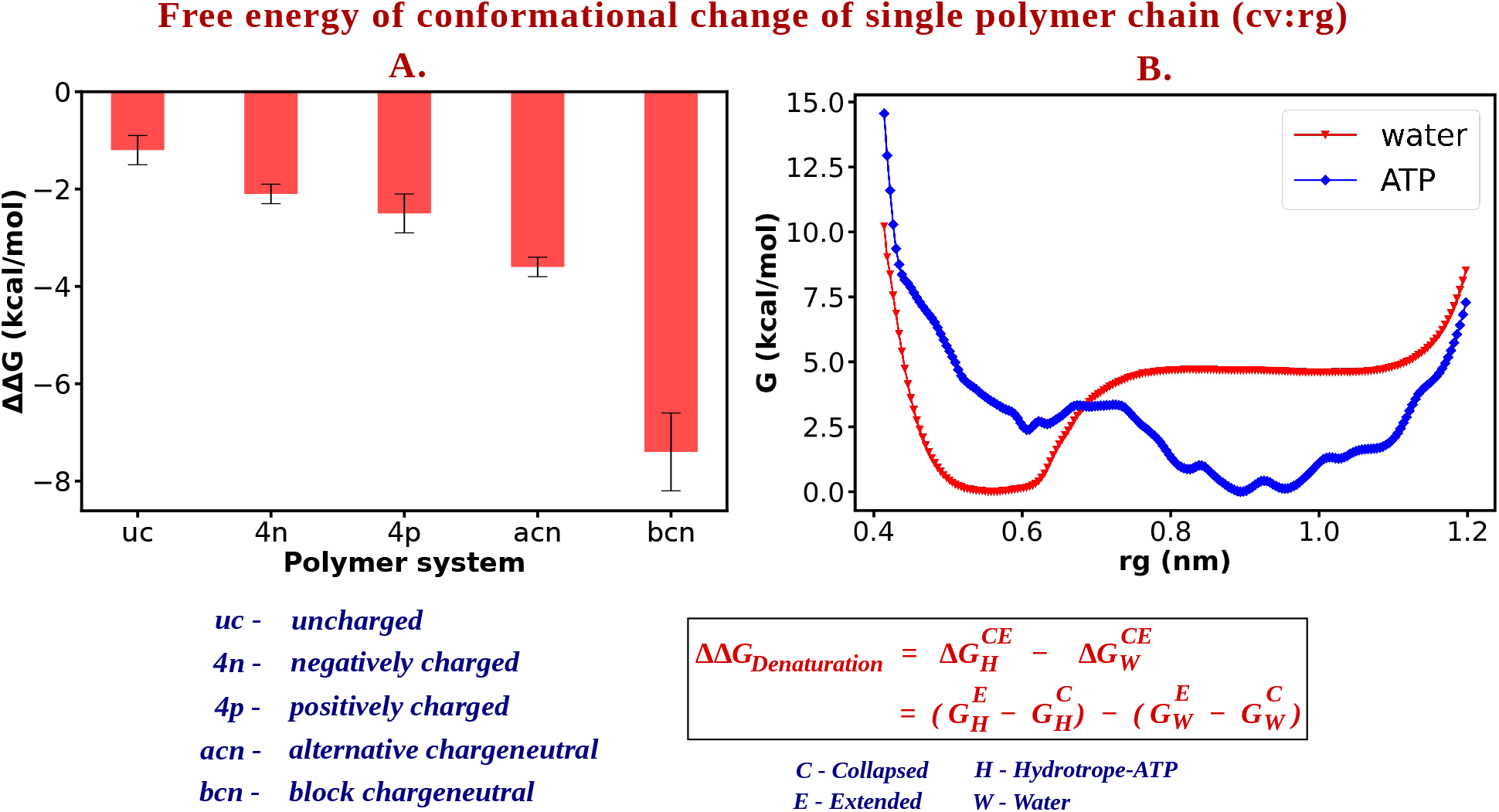
A. ΔΔG values along with the corresponding error bars for all the five polymer systems- (i) uncharged (ii) negatively charged, (iii) positively charged, (iv) alternative charge-neutral and (v) block chagre-neutral. B. Free energy profile of single block charge-neutral polymer in neat water (red) and in 0.6M aqueous ATP solution (blue)

### A preferential interaction based perspective of ATP’s action

The investigations have till now indicated that ATP has the intrinsic ability of both unfolding a single polymer chain and sollubilizing a multi-chain aggregate, a hall-mark of a hydrotrope. In the process, we have made an important observation: the presence of charge in the macromolecule reinforces the ATP’s efficiency in modulating the conformational landscape of single polymer chain towards its extended conformation. The question now arises, what is the mechanistic origin of ATP’s ability in these features ? Towards this end, we first quantitatively characterise ATP’s interaction with the macromolecules via certain key metrics.

A relevant parameter that is often employed in this regard and related work^19,33^ is the preferential binding coefficient (Γ) of ATP with the polymer surface over solvent water. Γ^16,17^ is defined as

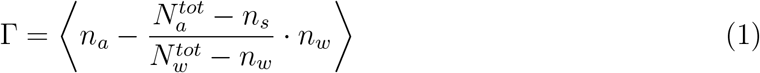

where *n_a_* represents the number of ATP molecules bound to the polymer and *N_a_^tot^* is the total number of ATP present in the system. On the other hand, *n_w_* stands for the number of water molecules bound to the polymer and *N_w_^tot^* is the total number of water molecules in the system. Γ quantitatively denotes the excess of ATP (referred as a) in the polymer solvation shell in comparison to its average concentration in the solution. Considering the rich chemistry involved in ATP’s structure, we measure Γ for three different parts of ATP i.e. triphosphate group (‘PG’), pentose sugar moiety (’sugar’) and the heterocyclic base part (‘base’). We have considered the central oxygen atom of water for our Γ calculation. In order to calculate the preferential binding coefficients (Γ) we carry out five independent simulations of 20 ns each by restraining the position of the polymers in collapsed (Rg: 0.45nm) or extended state (Rg: 1.0nm) for the single polymer for each aforementioned cases (5 different types of polymeric system). For attaining a better statistics, all the values of Γ have been averaged over 5 independent trajectories corresponding to the two conformations (collapsed (C) and extended (E)) of each of the different kinds of polymer chain. Γ > 0 would provide a measure for the preferential binding of ATP to the polymer surface, while a negative value of Γ indicates the depletion of ATP molecule from the polymer surface. Figure 6A represents the profile of Γ, for three different chemical moieties of ATP, as a function of distance from a charge-neutral (block charge-neutral) polymer chain in its two prominent conformations, namely collapsed and extended conformation in 0.6 M aqueous solution of ATP. Figure S4 represents the same for all five types of polymers investigated in the current work. Together, we find that Γ > 0 for all cases, indicating that ATP strongly binds to the polymer irrespective of its diverse coulombic nature. In the same figure, the representative snapshots of collapsed and extended conformations in a solution of ATP further illustrate the scenario. A closer inspection of the profiles in figure 6A indicates that, for a given polymer-ATP distance, the extended conformation has a large value of Γ than that of collapsed conformation, i.e. ΔΓ= Γ_*E*_−Γ_*C*_ > 0. This result is in accordance with Wyman-Tanford theory^16,17^, that relatively larger value of Γ near the extended conformation than that near collapsed conformation would drive the conformational equilibrium more towards extended state in an aqueous ATP solution. These results predict that more ATP molecules would be binding with the extended state of the polymer than the collapsed form and thereby would favour denatured state of a polymer chain via increasing the hydrophilic interaction associated with this conformation (as more ATP molecules with hydrophilic part can surround the extended form than the folded conformation). Overall, the investigation confirms ATP’s direct interaction with the polymer surface as the key mechanism. These analysis support the earlier predictions^1,34–36^ about the clustering of hydrotrope ATP around the hydrophobic substances during solubilization and is consistent with the basis of preferential binding theory. ^16,17^

**Figure 6:**
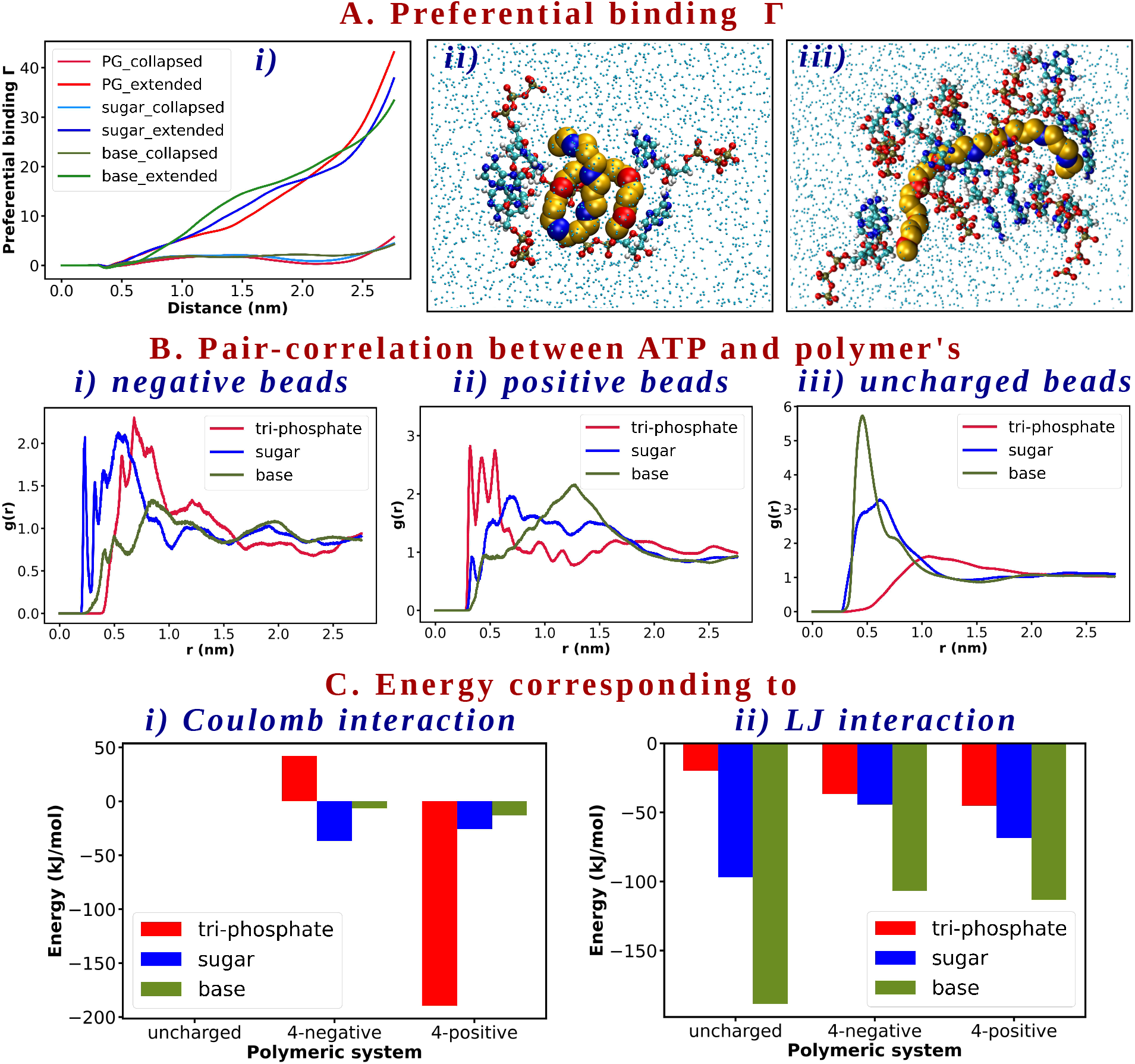
A. i) Distance profile of preferential binding coefficient (of block charge-neutral polymer), A.ii) and A.iii) are the representative snapshots for collapsed and extended state polymer (block charge-neutral) respectively showing more ATP is interacting with the extended conformer. B. RDF plot of different parts of ATP i.e. tri-phosphate (red), sugar (blue) and base (olive green) with respect to i) negative beads (of alternative charge-neutral polymer), positive beads (of block charge-neutral polymer) and uncharged beads (of purely uncharged polymer) C. Bar plot representation of i) Coulomb and ii) LJ interaction for the systems of single uncharged, negatively charged (4-negative) and positively charged (4-positive) polymers corresponding to the interaction of three different parts of ATP i.e. tri-phosphate (red), sugar(blue) and base (olive green)

The co-existence of positively charged, negatively charged and uncharged beads in a charge-neutral polymer provides an opportunity for jointly comparing the relative binding propensities of three different parts of ATP i.e. triphosphate group (‘PG’), pentose sugar moiety (‘sugar’) and the heterocyclic base part (‘base’) with the polymer surface. Figure 6B presents the radial distribution functions of different parts of ATP with the different types of beads of the charge-neutral polymer. We find that the sugar part of ATP is most strongly interacting with the negative beads. An atomistic analysis of g(r) (see figure S5) suggests that the CHOH group of sugar sets up the key interaction, as shown by the comparison of g(r) between different atoms of the sugar base and the polymer chain. On the other hand, it is the negatively charged phosphate group that interacts the most with the positively charged beads of the polymer. Finally, the radial distribution function in figure 6B suggests that the uncharged beads of polymers are predominantly interacting with the aromatic base part of ATP. This occurs mostly via purely hydrophobic interaction, in which the hydrophobic part of ATP (aromatic nitrogenous base) is selectively interacting with the hydrophobic uncharged polymer beads. The trend is common across all types of polymers (see figure S6 A,B and C).

The respective contribution of the interactions of different parts of ATP selectively with different types of beads of our model polymer is justified by the relative energies corresponding to each interaction between different parts of solubilizer (ATP) and solubilizate (hydrophobic polymer here). Here we have calculated both the coulombic and LJ interaction energy contribution during the process of solubilization. For this calculation we have used the trajectory produced after umbrella sampling simulation of the extended states (since ATP is interacting efficiently with extended conformer of the polymer) of the single polymer chain. Figure 6C compares the contributions of Coulomb and LJ interactions between different structural units of ATP with that of the different bead types of the charge-neutral polymer. We find that apolar base is mostly interacting with the polymer’s uncharged beads via hydrophobic interaction. Thus for base, the interaction is grossly contributed by LJ interaction. Whereas the coulomb interaction (electrostatic attraction) is mainly adopted between tetra-negatively charged tri-phosphate group and the positively charged beads of the polymer. On the other hand, for interaction involving sugar part of ATP, both the LJ and coulomb interaction come into play.

In summary, the depiction of preferential binding co-efficient of ATP relative to water establishes the mechanism of ATP’s hydrotropy via confirming direct and strong interaction of ATP with hydrophobic entities. The analysis of pair-correlation and interaction energy further informs about the mode of selective interaction of different parts of ATP with diverse polymer beads. The analysis is in line with the previous hypothesis drawn in Kang et al^9^ and Pal et al’s^10^ investigations about the key interactions driving the process and more importantly emphasises that the the ATP’s role get reinforced in presence of charge.

### Charge-reinforced ATP interaction drives Sollubilization

Overall the previous analysis underscores three key interactions behind ATP’s hydrotropic role-1) Favourable dispersion interaction between negative beads and ATP’s sugar part (mainly CHOH groups), 2) electrostatic interaction between positive beads and negatively charged phosphate group of ATP and 3) hydrophobic interaction between uncharged beads and hydrophobic aromatic base part of ATP. Thus in a system which would have uncharged, negative and positive beads coexisting (such as in a charge-neutral polymer), all these three interactions are expected to synergistically assist ATP to denature the polymer and also to solubilize polymeric aggregate more efficiently. The precedent analysis also prompts us to infer that ATP would behave as the most efficient hydrotrope for a charge-neutral system consisting three types of beads, compared to the other systems.

To test our hypothesis, we designed a series of simulation of the aggregation of charge-neutral polymeric systems in various aqueous ATP solution (simulation protocol and the concentration of ATP are same as employed for aggregation of uncharged polymer). Figure 7A compares the representative configuration of equilibrated aggregates of charge-neutral polymer in neat water and in 1 M aqueous ATP solution. In all cases, the simulations were initiated with different copies of polymers (5,10 and 20) randomly dispersed in solution. We find that while in neat water, the aggregation process of charge-neutral polymers results in a single aggregate for all investigated systems, the extent of aggregation is significantly diminished in 1 M ATP solution: either the chains remain mutually segregated in lower polymer copy numbers or remain solubilized in multiple smaller-sized aggregates. For a pertinent comparison with the uncharged hydrophobic polymer, we refer to figure 4, where it was observed that although the uncharged polymers avoid the formation of a single large aggregate in ATP solution, the size of the aggregates formed of uncharged polymers in 1 M ATP is still relatively larger than that in charge-neutral polymer in same ATP concentration. For a quantitative characterisation of this charge-reinforced hydrotropic effect, we analyse the cluster size distributions in ‘uncharged’, ‘alternatively charge-neutral’ and ‘block charge-neutral’ polymers in near water and 1 M ATP. Figure 7B compares the number of clusters in these three cases in neat water and 1 M ATP solution. It is evident, while the number of clusters increases from 1 (aggregate constituted of all polymers in solution) in neat water to multiple copies of smaller-sized clusters (than a single aggregate) in 1 M ATP in all cases, the size of the aggregates is relatively smaller in both charge-neutral systems. As shown in figure 7B, at 1M ATP, in the polymer system with 5 copies of uncharged polymers, two smaller aggregates are formed consisting of 2 and 3 polymer chains in each of them. On the other hand, for both the charge-neutral polymer systems (alternative and block charge-neutral) the number of chains in cluster is 1 in 1M ATP, (represented by blue and green bar with the number ‘1’ at top of each bar), echoing ATP’s ability in keeping the chains mutually isolated. A very similar picture emerges in 10 or 20 copies as well, as evident from the cluster size analysis in figure 7B.

**Figure 7:**
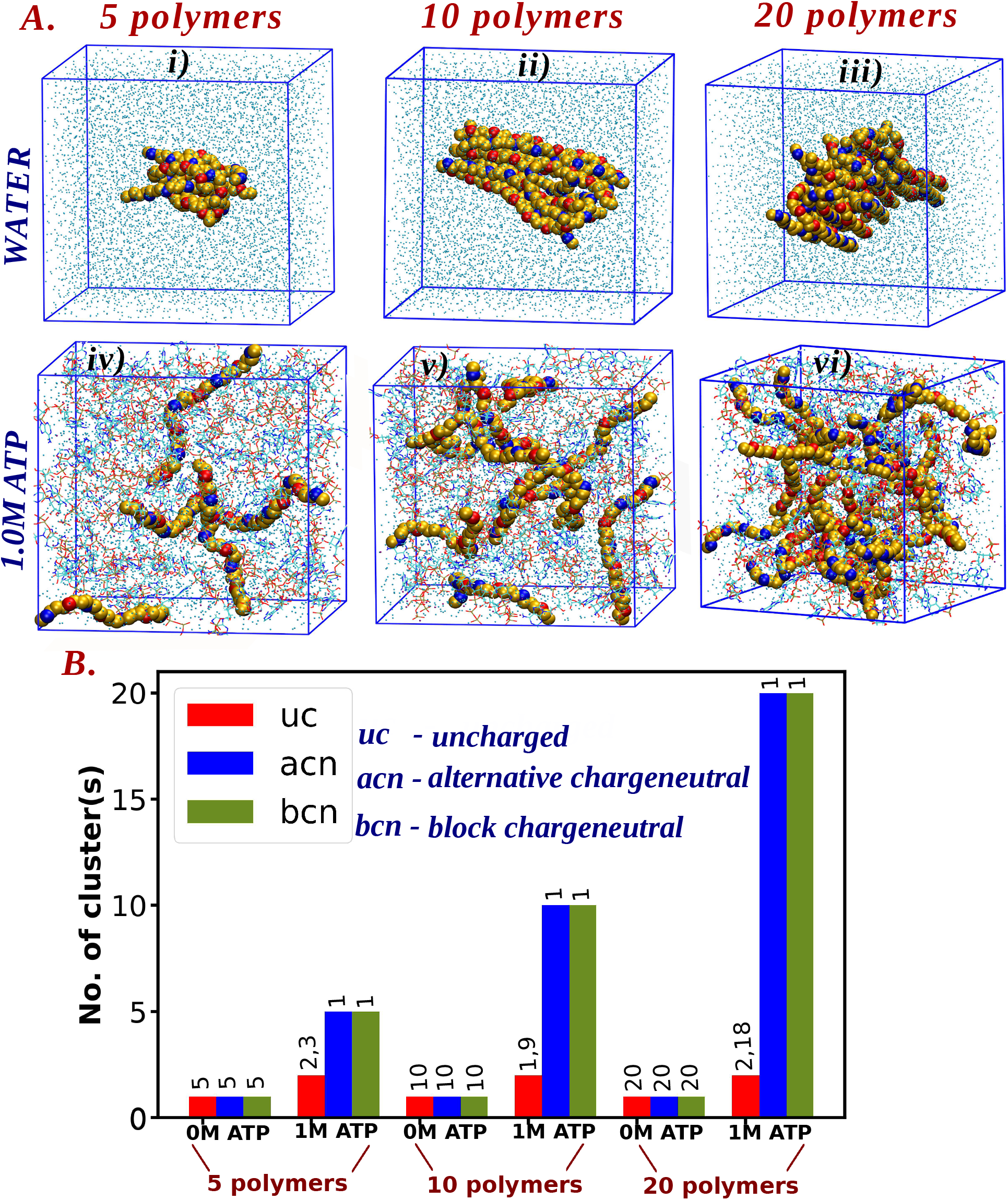
A. formation of single large aggregate (consisting of all the polymers present in the system) for the case of i) 5, ii) 10 and iii) 20 charge-neutral polymers in absence of ATP i.e in neat water medium and figure iv-vi) represents ATP induced segregation for those polymer systems of 5, 10 and 20 charge-neutral chains in 1 M ATP. B. Cluster size distribution for the systems with i) 5, ii) 10, and iii) 20 uncharged and both the charge-neutral (alternative and block charge-neutral) polymers in 0 M and 1 M ATP concentrations.

On a similar note, the cluster-analysis in figure S7 demonstrates that the threshold concentration of ATP required to dissolve aggregates of charge-neutral polymers is significantly lower than that of purely uncharged polymeric aggregates. The complete solubilization of the aggregate, formed of hydrophobic uncharged polymers, into segregated individual chains was not realised even in 1 M aqueous solution of ATP. On the contrary, 0.4M aqueous ATP solution was found to be sufficient to efficiently dissolve the aggregate of both the charge-neutral polymers into single individual monomers. Specifically, for both the charge-neutral systems, each of the polymer chains remained mutually segregated upon the treatment of only 0.4M ATP. This signifies that the hydrotropic or sollubilizing effect of ATP is relatively stronger towards charge-neutral polymers than purely uncharged one.

For a quantitative characterisation of effect of charges, we computed the free energy of self-assembly of a pair of each of the polymers in neat water and in 0.6M M ATP concentration. Figure 8B depicts the computed free energy profiles as a function of distance of centre of mass between two charge-neutral polymer chains, while Figure S8 put together similar free energy profiles for all sets of polymers investigated in the current work. In all cases, the qualitative trend of ATP-induced destabilisation of self-assembled aggregate holds good. However, the extent of destabilisation is most prominent in charge-neutral polymer, followed by net charged polymer. As a quantitative comparison, we compute the relative change in free energy (ΔΔ*G*) of aggregation in ATP solution over neat water across different polymer types. A negative value of ΔΔ*G* indicates that the aggregate is free energetically more destabilised in ATP solution that that in neat water. Figure 8A compares the value of ΔΔ*G* of aggregation for all five types of polymers. The extent of ATP-induced destabilisation of aggregation is maximum in case of charge-neutral polymer, which echoes the observation of our investigation of aggregation process.

**Figure 8:**
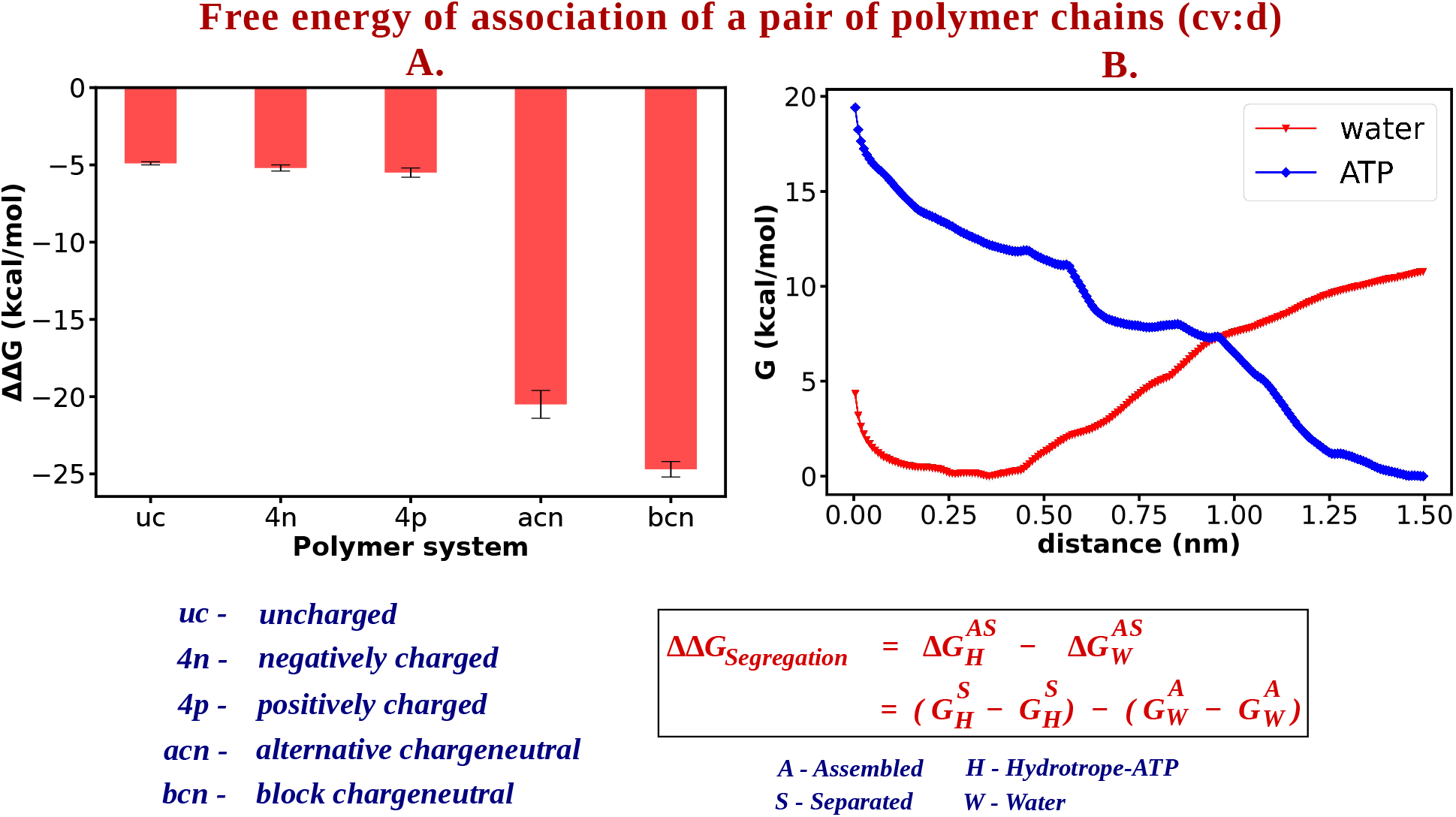
A. ΔΔG values for all the five polymer systems- (i) uncharged (ii) negatively charged, (iii) positively charged, (iv) alternative charge-neutral and (v) block chagre-neutral. B. Free energy profile of single block charge-neutral polymer in neat water (red) and in 0.6M aqueous ATP solution (blue)

### Self-aggregation of ATP makes it an efficient hydrotrope

As an intriguing observation from our investigation, we noted that ATP molecules were self-aggregating in the aqueous medium. Figure 9A depicts a representative snapshot of ATP molecules in 0.08 M Aqueous ATP media where the clustering of ATP is clearly evident. The possibility of ATP’s self-aggregation makes sense from a chemical perspective. There are three possible interactions associated with the process of aggregation of ATP molecules- 1) *π*−*π* interaction between the aromatic rings of the adjacent molecules, 2) anion−*π* interaction between the aromatic base ring and the negatively charged oxygen moiety of the triphosphate group in ATP and additionally there is a chance of 3) hydrogen bonding interaction between the sugar OH group and the triphosphate oxygen anion. All of these interactions likely co-operatively act together to bring ATP molecules closer to each other.

**Figure 9:**
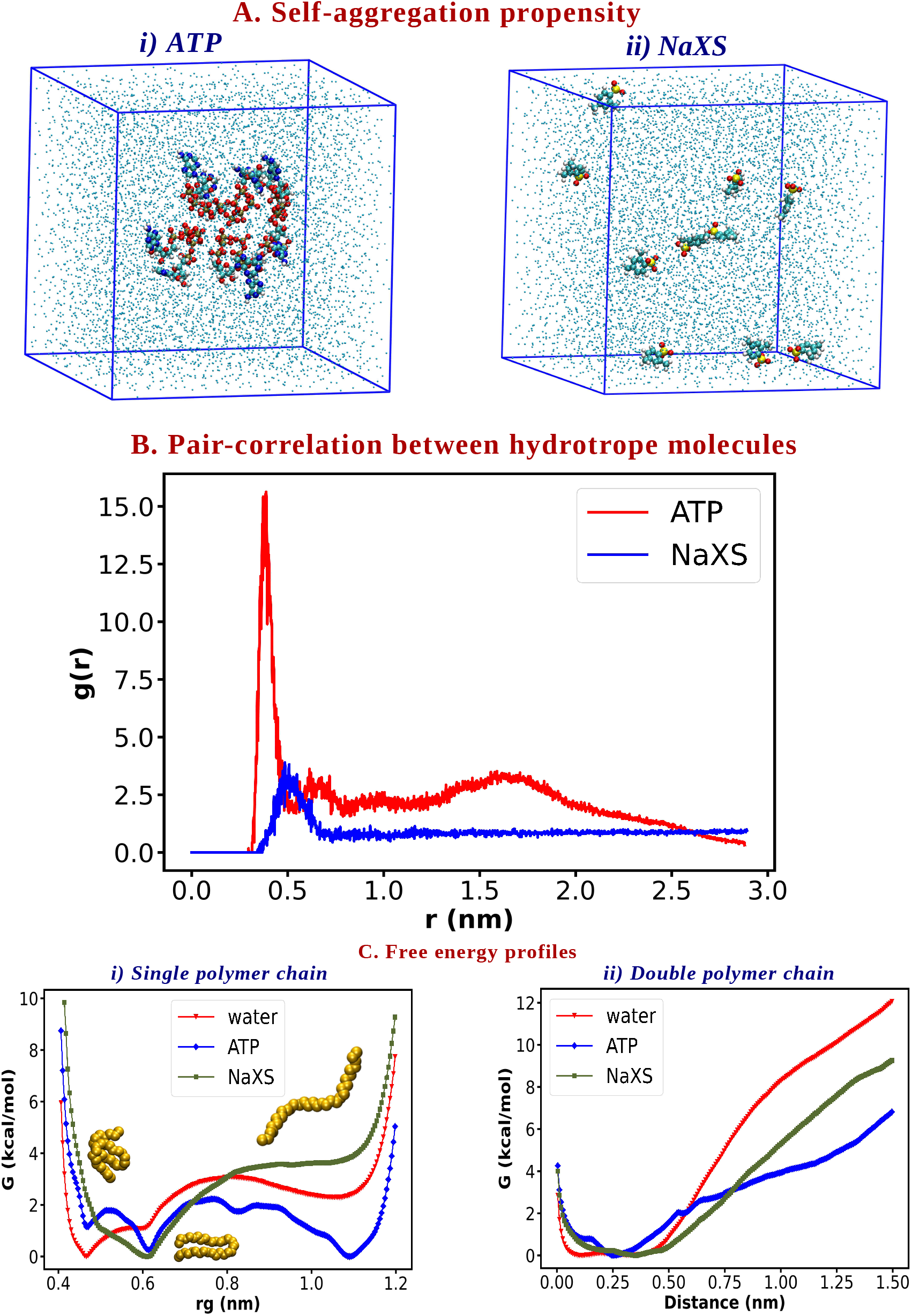
A. Snapshot showing that self-assembly of i) ATP in neat water but the same is not observed in case of ii) NaXS. B. A representation of pair-correlation between the hydrotrope molecules ATP (red) and NaXS (blue) C. Free energy profiles comparing the energy landscapes in neat water, 0.6M ATP solution and 0.6M NaXS solution for (i) single uncharged polymer and (ii) double uncharged polymers.

Interestingly, experiments comparing other ‘chemical hydrotropes’ NaXS with ATP^1^ have shown that significantly lower concentration of ATP is sufficient to do the job even more efficiently than NaXS. We posited that this phenomenon of self-aggregation in ATP can be identified as one of the major reasons towards the efficiency of ATP as a denaturant and a solubilizer. Since ATP has an inherent tendency to self-aggregate, the local concentration of ATP increases near polymer surface, which might act as a pre-requisite for creating an effectively higher ATP concentration around the hydrophobic solute. On the other hand, for the other hydrotrope, which lacks the property of self-aggregation, one needs to maintain comparatively higher concentration of the same to reach the concentration actually required for its proper action.

Accordingly, we carried out additional control simulations with a typical chemical hydrotrope NaXS (the same as used by Patel et al.^1^) Interestingly we encountered that unlike ATP, NaXS does not self-aggregate (figure 9B). ATP’s relatively higher self-aggregation propensity is also confirmed by a comparison of radial distribution function. We believe that it is mainly because of the presence of negatively charged moiety (just adjacent to the aromatic ring) in NaXS molecule, the closer approach of two NaXS molecules can not be permitted (due to severe electrostatic repulsion among them), precluding their self-aggregation. On the other hand, since the negatively charged triphosphate group in ATP is remotely placed from the aromatic ring, the repulsion among them could not resist the *π*−*π* interaction between the aromatic rings. For a conclusive evidence of ATP’s superior unfolding and sollubilizing ability over NaXS, we compare the free energetics of conformational landscape and aggregation propensity between neat water, 0.6 M aqueous ATP and 0.6 M aqueous NaXS solution (figure 9C). The free energy profiles clearly show that ATP will be significantly more efficient in unfolding a polymer chain and solubilizing an aggregate than NaXS at the same concentration, indicating ATP’s self-aggregating propensity key to its superior efficacy over NaXS.

## Conclusion

Considering its role as cellular energy currency, ATP has always secured special position in biology. The recent discovery of ATP’s additional hydrotropic ability in multiple crucial biological aspects (maintaining the high concentration of proteins soluble in eye lens,^5^ influencing liquid-liquid phase separation,^1^ inhibiting neurogenerative diseases^10^ etc.) have further strengthened ATP’s powerful contribution in biology. These investigations in turn have demanded the evolution of detailed atomistic mechanism behind ATP’s hydrotropy and the associated driving forces. Although “hydrotropy” is a much older concept, the actual mechanism around this has long been a debate. In our study we have tried to establish the mechanistic details of ATP’s hydrotropy based on its denaturation and segregation ability.

In summary, the current article has investigated the possible mechanism of recently discovered role of ATP as a biological hydrotrope via a bottom-up approach. In stead of scrambling with complex interplay of protein condensates, here we computationally designed simple prototypical polymers and explored the action of differently concentrated ATP solution on it via MD simulation. Specifically, our results combining equilibrium and free energy simulations bring out dual roles of ATP: 1) ATP denatures the native globular conformation of a single polymer chain to its extended form and 2) it disrupts aggregation process in neat water. An interesting result that emerged from current simulations is that the presence of charges in the solute would reinforce ATP’s hydrotropic role. Especially, ATP was found to prevent the aggregation process of a charge-neutral polymer more efficiently than a hydrophobic polymer. A molecular interaction based analysis points to ATP’s preferential binding to the polymer surface (relative to water) via dispersion and coulombic interaction with different structural units as the key to ATP’s denaturing and sollubilizing role. Overall, the prediction that the presence of charge in macromolecule would bolster ATP’s hydrotropic role is in sync with prior experimental observation ^1^ of solubilisation of intrinsically disordered proteins (which are generally very rich in charged amino acids) in aqueous ATP solution. The charge-reinforced effect of ATP, as found in the current simulation, is further supported by recent experimental report of Kang et al^9^ and computer simulation of Pal et al^10^ which investigated ATP’s hydrotropy on FUS and A*β*16-22 respectively and enumerated different interactions acting during disaggregation and most of those were majorly contributed from electrostatics.

As an interesting observation from our simulation, we noticed that the key to ATP being more effective hydrotrope than a ‘chemical hydrotrope’ NaXS lies in the former’s (i.e. ATP) inherent self-assembling propensity, otherwise absent in NaXS. We provide circumstantial evidence that the extent of self-aggregation of ATP regulates the concentration of ATP within the cells, which might explain its occurrences in cell at higher concentration ^1^ than that is required for its role as energy currency. Our simulations suggest that the self-assembling propensity can cause an increase in local crowding of ATP molecules, which would enhance ATP’s binding propensity with the macromolecule, thereby boosting ATP’s hydrotropic effect. The picture emerging from the current simulation results is qualitatively consistent with the proposal of dynamic clustering of ATP around solute particles, previously put forward by past theory^34–36^ and experiment.^1^ Since the formation of clusters of hydrotrope over solubilizate lies in the heart of hydrotropic activity, this additional factor (ATP’s self-assembly) enhances ATP’s propensity of accumulation and thereby intensifies the degree of dynamic clustering.

Overall this work provides an intuition towards the importance of the presence of ATP to make the proteins soluble within cells. It will be worth exploring if the predictions made based on simple macromolecule can be translated to realistic protein system, which we plan to pursue in a future work. Since ATP can enormously slow down the process of aggregation, its presence in solution might possibly help an improved investigation of intrinsically disordered protein (which otherwise undergoes rapid fluctuation). Since ATP helps in disaggregation, we have our future plan to test ATP’s effect in prion like protein-aggregation.^37^

## Supplemental Information

All supplemental figures described in the article (PDF)

## Acknowledgements

This work was supported by computing resources obtained from shared facility of TIFR Centre for Interdisciplinary Sciences, India. We acknowledge support of the Department of Atomic Energy, Government of India, under Project Identification No. RTI 4007. JM acknowledges Ramanujan Fellowship and Core Research grants provided by the Department of Science and Technology (DST) of India (CRG/2019/001219).

---

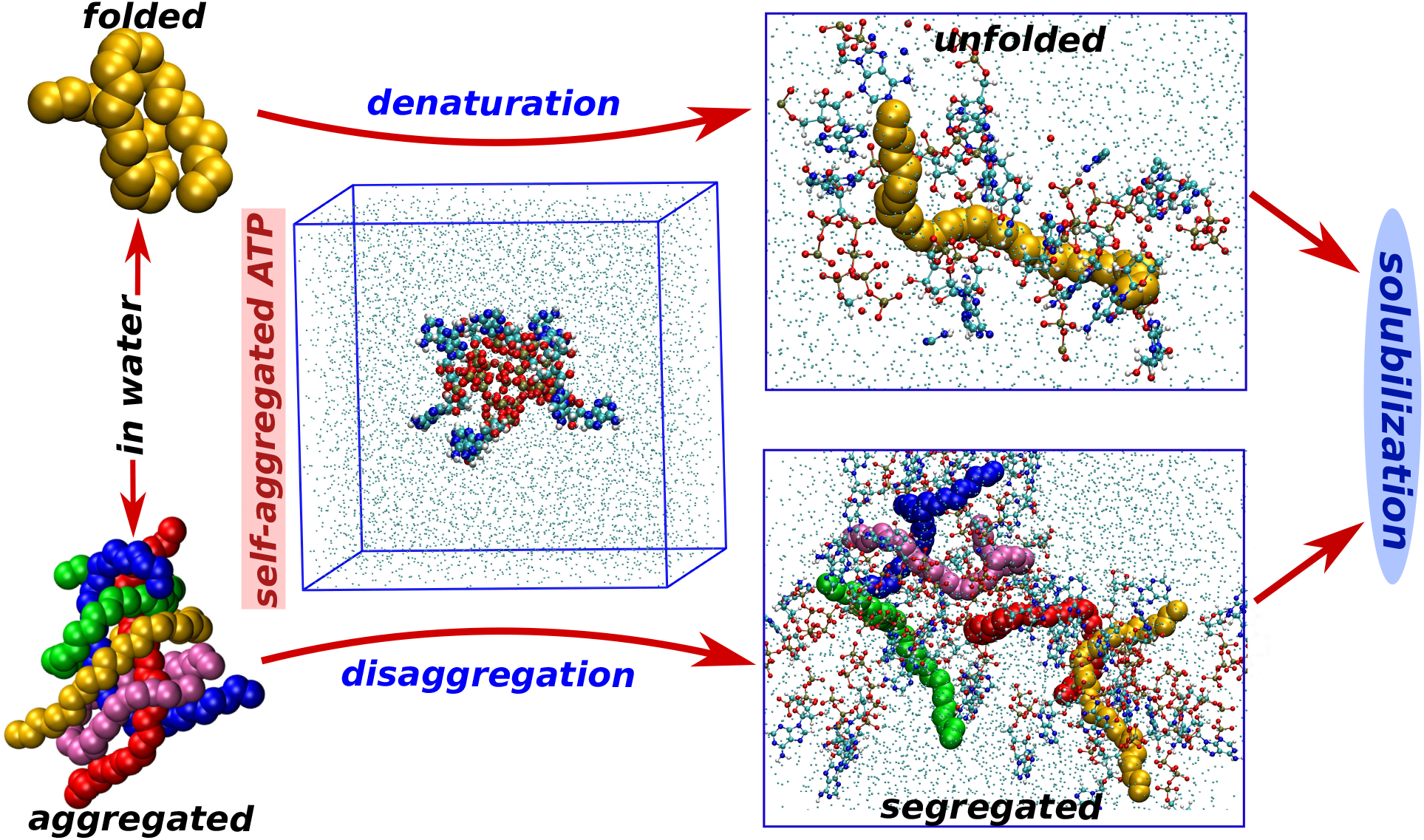

## References

(1) Patel, A.; Malinovska, L.; Saha, S.; Wang, J.; Alberti, S.; Krishnan, Y.; Hyman, A. A. ATP as a biological hydrotrope. Science 2017, 356, 753–756.

(2) Brangwynne, C. P.; Mitchison, T. J.; Hyman, A. A. Active liquid-like behavior of nucleoli determines their size and shape in Xenopus laevis oocytes. Proceedings of the National Academy of Sciences 2011, 108, 4334–4339.

(3) Jain, S.; Wheeler, J. R.; Walters, R. W.; Agrawal, A.; Barsic, A.; Parker, R. ATPase-Modulated Stress Granules Contain a Diverse Proteome and Substructure. Cell 2016, 164, 487–498.

(4) Hyman, A. A.; Weber, C. A.; Jülicher, F. Liquid-Liquid Phase Separation in Biology. Annual Review of Cell and Developmental Biology 2014, 30, 39–58.

(5) He, Y.; Kang, J.; Song, J. ATP antagonizes the crowding-induced destabilization of the human eye-lens protein *γ* S-crystallin. Biochemical and Biophysical Research Communications 2020, 526, 1112–1117.

(6) Hayes, M. H.; Peuchen, E. H.; Dovichi, N.; Weeks, D. L. Dual roles for ATP in the regulation of phase separated protein aggregates in Xenopus oocyte nucleoli. eLife 2018, 7, e35224.

(7) Sridharan, S.; Kurzawa, N.; Werner, T.; Gunther, I.; Helm, D.; Huber, W.; Bantscheff, M.; Savitski, M. M. Proteome-wide solubility and thermal stability profiling reveals distinct regulatory roles for ATP. Nature Comm. 2019, 10, e35224.

(8) Neuberg, C. In Biochemical Reductions at the Expense of Sugars; Pigm, W., Wolfro, M., Eds.; Advances in Carbohydrate Chemistry; Academic Press, 1949; Vol. 4; pp 75–117.

(9) Kang, J.; Lim, L.; Lu, Y.; Song, J. A unified mechanism for LLPS of ALS/FTLD-causing FUS as well as its modulation by ATP and oligonucleic acids. PLOS Biology 2019, 17, e3000327.

(10) Pal, S.; Paul, S. ATP Controls the Aggregation of A*β*16-22 Peptides. J. Phys. Chem. B 2019, 124, 210–223.

(11) McKee, R. H. Use of Hydrotropic Solutions in Industry. Industrial & Engineering Chemistry 1946, 38, 382–384.

(12) Winsor, P. A. Hydrotropy, solubilisation and related emulsification processes. Transactions of the Faraday Society 1948, 44, 376.

(13) Ueda, S. The Mechanism of Solubilization of Water Insoluble Substances with Sodium Benzoate Derivatives. I. The Interaction between Water Insoluble Substances and Sodium Benzoate Derivatives in Aqueous Solution. Chemical and Pharmaceutical Bulletin 1966, 14, 22–29.

(14) Kumar, V. S.; Raja, C.; Jayakumar, C. A review on Solubility Enhancement Using Hydrotropic Phenomena. Int. J. Pharm. Pharm. Sci. 2014, 6, 1–7.

(15) Balasubramanian, D.; Srinivas, V.; Gaikar, V. G.; Sharma, M. M. Aggregation behavior of hydrotropic compounds in aqueous solution. The J. Phys. Chem. 1989, 93, 3865–3870.

(16) Wyman, J. Linked Functions and Reciprocal Effects in Hemoglobin: A Second Look. Adv. Protein Chem. 1964, 19, 223–286.

(17) Tanford, C. Extension of the theory of linked functions to incorporate the effects of protein hydration. J. Mol. Biol. 1969, 39, 539–544.

(18) Torrie, G.; Valleau, J. Nonphysical sampling distributions in Monte Carlo free-energy estimation: Umbrella sampling. Journal of Computational Physics 1977, 23, 187 – 199.

(19) Mondal, J.; Stirnemann, G.; Berne, B. J. When Does TrimethylamineN-Oxide Fold a Polymer Chain and Urea Unfold It? J. Phys. Chem. B 2013, 117, 8723–8732.

(20) MacKerell, A. D. et al. All-Atom Empirical Potential for Molecular Modeling and Dynamics Studies of Proteins. J. Phys. Chem. B 1998, 102, 3586–3616, PMID: 24889800.

(21) Hart, K.; Foloppe, N.; Baker, C. M.; Denning, E. J.; Nilsson, L.; MacKerell, A. D. Optimization of the CHARMM Additive Force Field for DNA: Improved Treatment of the BI/BII Conformational Equilibrium. J. Chem. Theory Comput. 2012, 8, 348–362.

(22) Vanommeslaeghe, K.; Hatcher, E.; Acharya, C.; Kundu, S.; Zhong, S.; Shim, J.; Darian, E.; Guvench, O.; Lopes, P.; Vorobyov, I.; Mackerell Jr., A. D. CHARMM general force field: A force field for drug-like molecules compatible with the CHARMM all-atom additive biological force fields. J. Comp. Chem 2010, 31, 671–690.

(23) Huang, L.; Roux, B. Automated Force Field Parameterization for Nonpolarizable and Polarizable Atomic Models Based on Ab Initio Target Data. J. Chem. Theory Comput. 2013, 9, 3543–3556.

(24) Parrinello, M.; Rahman, A. Polymorphic transitions in single crystals: A new molecular dynamics method. J. Appl. Phys. 1981, 52, 7182–7190.

(25) Darden, T.; York, D.; Pedersen, L. Particle mesh Ewald: AnN·log(N) method for Ewald sums in large systems. J. Chem. Phys. 1993, 98, 10089–10092.

(26) Hess, B.; Bekker, H.; Berendsen, H. J. C.; Fraaije, J. G. E. M. LINCS: A linear constraint solver for molecular simulations. J. Comp. Chem 1997, 18, 1463–1472.

(27) Miyamoto, S.; Kollman, P. A. Settle: An analytical version of the SHAKE and RATTLE algorithm for rigid water models. J. Comp. Chem 1992, 13, 952–962.

(28) Abraham, M. J.; Murtola, T.; Schulz, R.; Pall, S.; Smith, J. C.; Hess, B.; Lindahl, E. GROMACS: High performance molecular simulations through multi-level parallelism from laptops to supercomputers. SoftwareX 2015, 1-2, 19–25.

(29) Hess, B.; Kutzner, C.; van der Spoel, D.; Lindahl, E. GROMACS 4: Algorithms for Highly Efficient, Load-Balanced, and Scalable Molecular Simulation. J. Chem. Theory Comput. 2008, 4, 435–447, PMID: 26620784.

(30) Bonomi, M.; Branduardi, D.; Bussi, G.; Camilloni, C.; Parrinello, M. PLUMED: A portable plugin for free-energy calculations with molecular dynamics. Comp. Phys. Comm. 2009, 180, 1961–1972.

(31) Tribello, G. A.; Bonomi, M.; Branduardi, D.; Camilloni, C.; Bussi, G. {PLUMED} 2: New feathers for an old bird. Comput. Phys. Comm. 2014, 185, 604 – 613.

(32) A. Grossfield, WHAM: the weighted histogram analysis method, version 2.0.9. http://membrane.urmc.rochester.edu/content/version-209, 2014.

(33) Mondal, J.; Halverson, D.; Li, I. T. S.; Stirnemann, G.; Walker, G. C.; Berne, B. J. How osmolytes influence hydrophobic polymer conformations: A unified view from experiment and theory. Proc. Natl. Acad. Sci. USA 2015, 112, 9270–9275.

(34) Booth, J. L.; Abbott, S.; Shimizu, S. Mechanism of Hydrophobic Drug Solubilization by Small Molecule Hydrotropes. J. Phys. Chem. B 2012, 116, 14915–14921.

(35) Booth, J. J.; Omar, M.; Abbott, S.; Shimizu, S. Hydrotrope accumulation around the drug: the driving force for solubilization and minimum hydrotrope concentration for nicotinamide and urea. Phys. Chem. Chem. Phys. 2015, 17, 8028–8037.

(36) Shimizu, S.; Matubayasi, N. Hydrotropy: Monomer and ÄìMicelle Equilibrium and Minimum Hydrotrope Concentration. J. Phys. Chem. B 2014, 118, 10515–10524, PMID: 25144510.

(37) Raimundo, A. F.; Ferreira, S.; Martins, I. C.; Menezes, R. Islet Amyloid Polypeptide: A Partner in Crime in the Pathology of Alzheimer’s Disease. Frontiers in Molecular Neuroscience 2020, 13, 35.

