## Supplemental Figures for "Mechanistic Insights on ATP’s role as Hydrotrope"

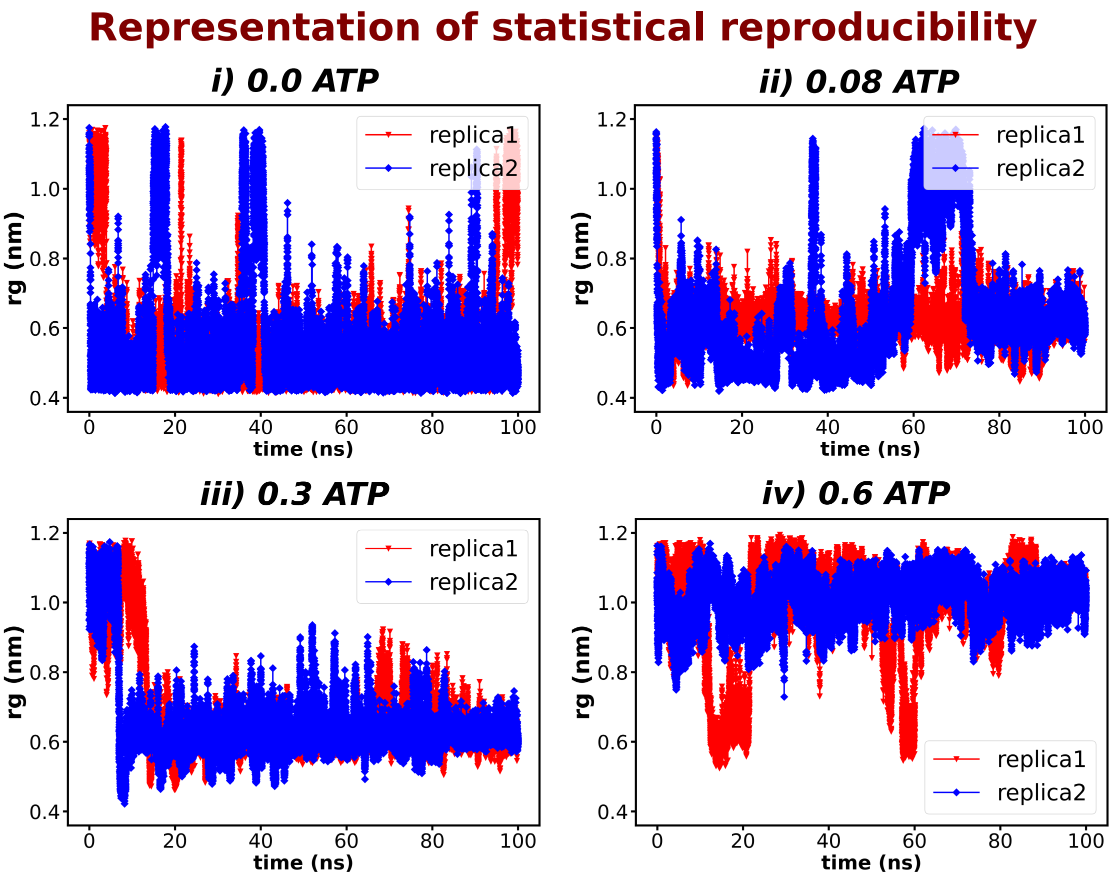


Figure S1: The time profile of $R_g$ of single uncharged polymer in 4 different ATP concentration 0.0M, 0.08M, 0.3M and 0.6M for two independent trajectories

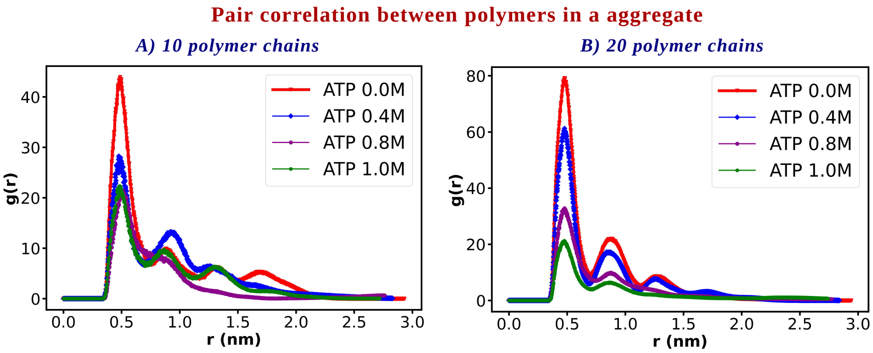


Figure S2: Pair-correlation between all the uncharged polymer chains involved in aggregate in 4 different concentrations of ATP 0.0M, 0.4M, 0.8M and 1M for two polymeric systems with i) 10 and ii) 20 uncharged chains

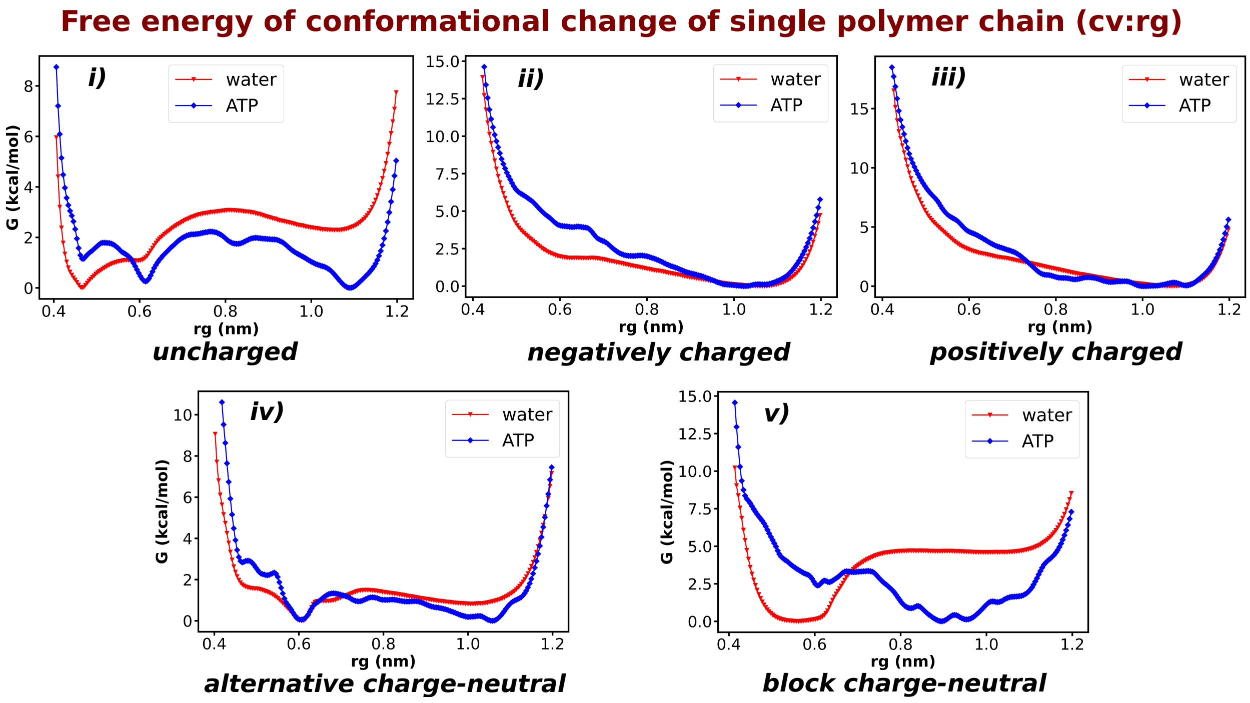


Figure S3: Free energy profile for the single polymer chain of all the five polymeric systems i) uncharged, ii) negatively charged, iii) positively charged, iv) alternative charge-neutral and v) block charge-neutral at 0.0M and 0.6M aqueous ATP solution

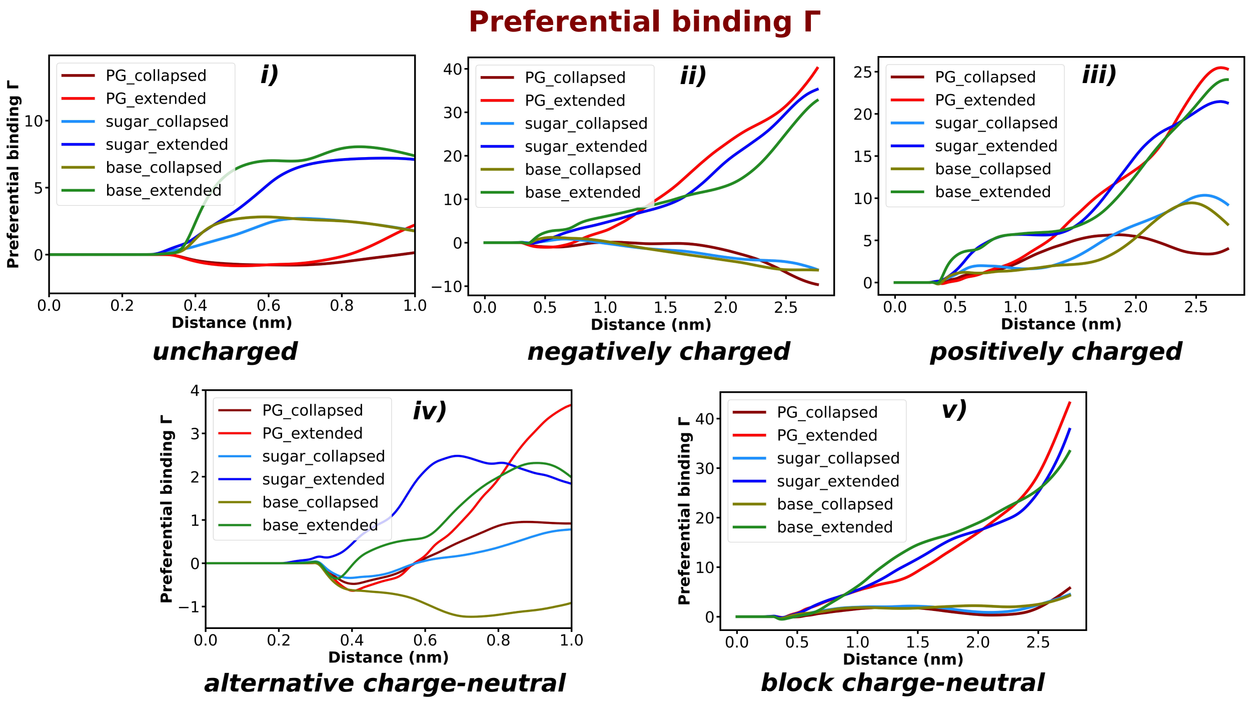


Figure S4: Distance profile of preferential binding co-efficient Γ with respect to three different parts of ATP i.e. triphosphate, sugar and base with respect to both the extended and collapsed conformation of each of the polymeric system i) uncharged, ii) negatively charged, iii) positively charged, iv) alternative charge-neutral and v) block charge-neutral

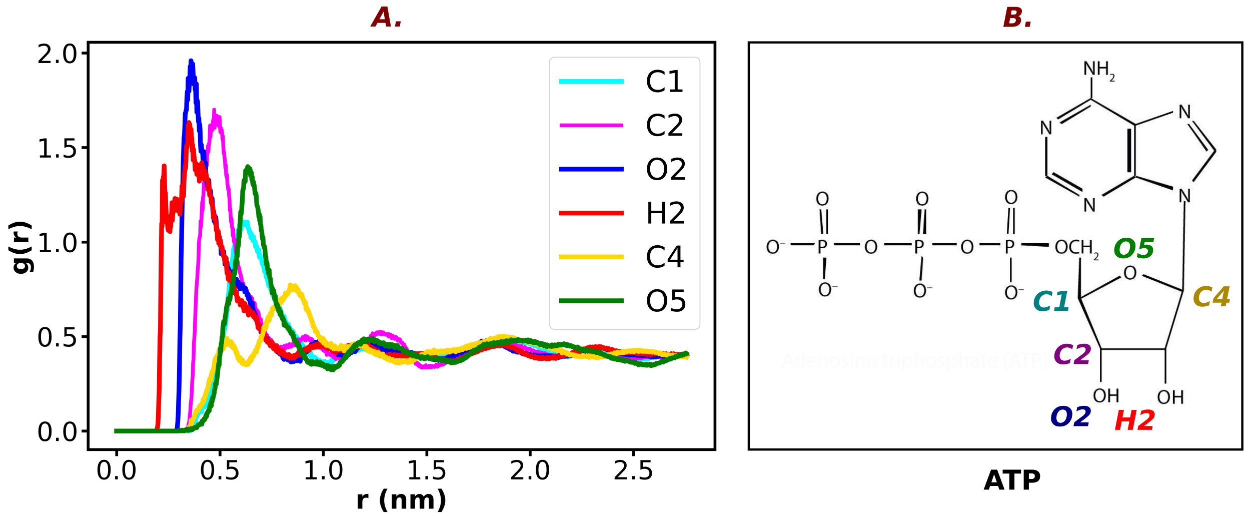


Figure S5: A. The pair-correlation of different atoms of the sugar group in ATP (shown in figure B) with respect to the negatively charged beads (of alternative charge-neutral polymer)

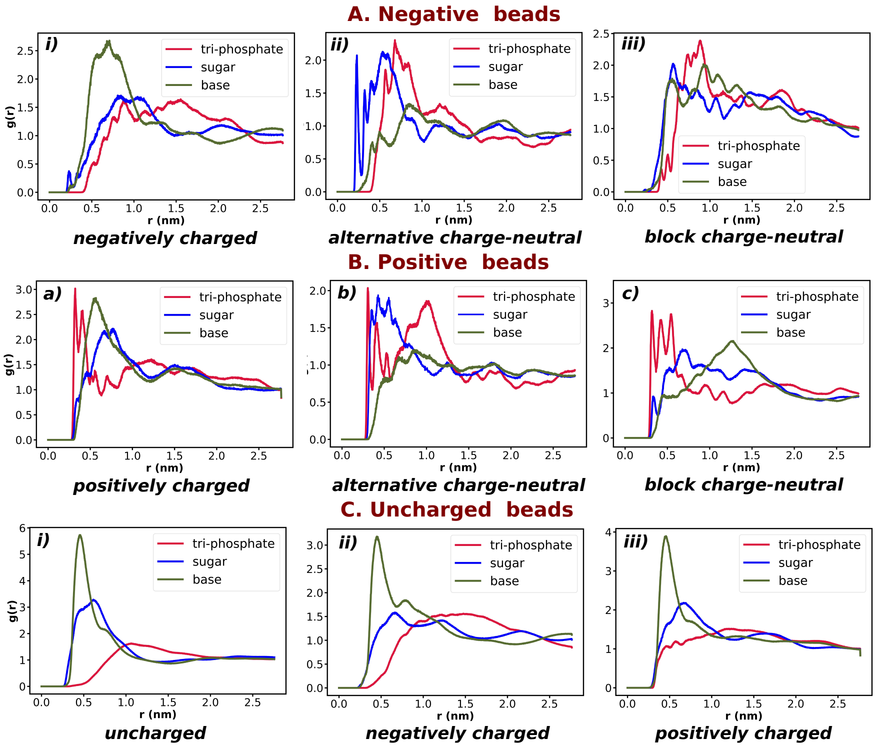


Figure S6: Pair correlation of different parts of ATP with different kinds of beads i) negative beads, ii) positive beads and iii) uncharged beads of the respective bead containing polymer systems

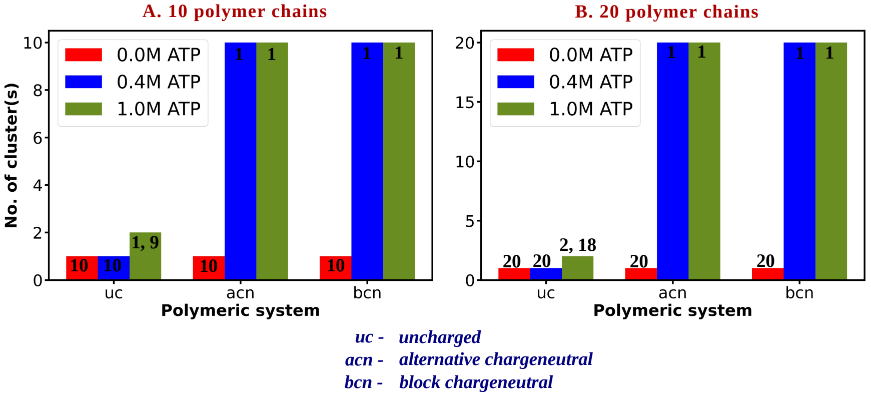


Figure S7: Bar plot representation of cluster-size distribution in A. 10 and B. 20 polymer systems at three different concentration of ATP for uncharged and both charge-neutral polymers (alternative and block charge-neutral) (The numbers have been shown corresponding to each bar represent the number of constituent polymers in each cluster)

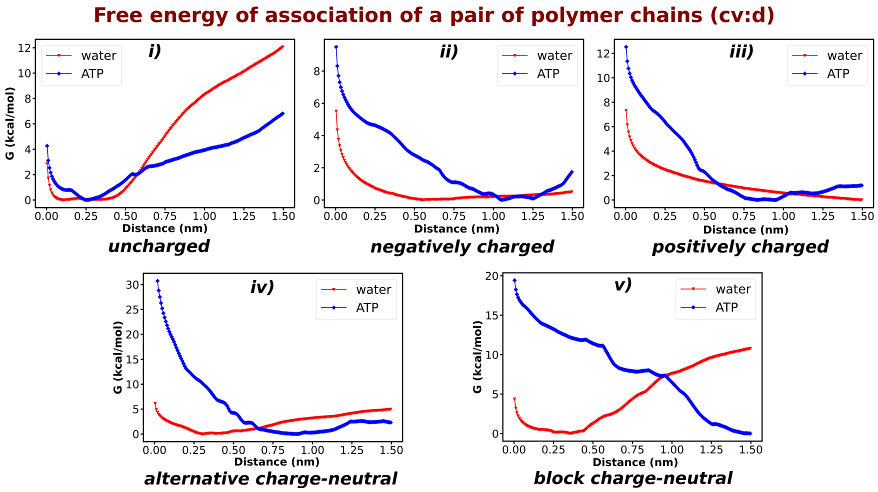


Figure S8: Free energy profile for association of a polymer chains of all the five polymeric systems i) uncharged, ii) negatively charged, iii) positively charged, iv) alternative charge-neutral and v) block charge-neutral at 0.0M and 0.6M aqueous ATP solution
